# *Hox*-dependent coordination of cardiac progenitor cell patterning and differentiation

**DOI:** 10.1101/2020.01.23.916981

**Authors:** Sonia Stefanovic, Brigitte Laforest, Jean-Pierre Desvignes, Fabienne Lescroart, Laurent Argiro, Corinne Maurel-Zaffran, David Salgado, Christopher de Bono, Kristijan Pazur, Magali Théveniau-Ruissy, Christophe Béroud, Michel Pucéat, Anthony Gavalas, Robert G. Kelly, Stéphane Zaffran

## Abstract

Perturbation of addition of second heart field (SHF) cardiac progenitor cells to the poles of the heart tube results in congenital heart defects (CHD). The transcriptional programs and upstream regulatory events operating in different subpopulations of the SHF remain unclear. Here, we profile the transcriptome and chromatin accessibility of anterior and posterior SHF sub-populations at genome-wide levels and demonstrate that Hoxb1 negatively regulates differentiation in the posterior SHF. Spatial mis-expression of *Hoxb1* in the anterior SHF results in hypoplastic right ventricle. Activation of *Hoxb1* in embryonic stem cells arrests cardiac differentiation, whereas *Hoxb1*-deficient embryos display premature cardiac differentiation. Moreover, ectopic differentiation in the posterior SHF of embryos lacking both *Hoxb1* and its paralog *Hoxa1* results in atrioventricular septal defects. Our results show that Hoxb1 plays a key role in patterning cardiac progenitor cells that contribute to both cardiac poles and provide new insights into the pathogenesis of CHD.

## INTRODUCTION

Heart morphogenesis and patterning require precise temporal differentiation of distinct cardiac progenitor populations that arise from two early sources of mesoderm progenitors, the first heart field (FHF) and the second heart field (SHF) (Buckingham et al., 2005). The FHF originates from the anterior splanchnic mesoderm and forms the cardiac crescent. The SHF is a progenitor population originating in the pharyngeal mesoderm that contributes to heart tube elongation through the progressive addition of cells from the dorsal pericardial wall to both poles of the forming heart. The SHF gives rise to right ventricular and outflow tract myocardium at the arterial pole, and to atrial myocardium including the dorsal mesenchymal protrusion (DMP) at the venous pole (Zaffran and Kelly, 2012). In the absence of SHF cell addition, impaired heart tube elongation and looping leads to early embryonic lethality, and perturbation of this process underlies a spectrum of common congenital heart defects (CHDs) (Prall et al., 2007; Cai et al., 2003). Lineage tracing analysis in mammals has revealed that the SHF is sub-divided into distinct anterior and posterior regions (aSHF and pSHF) (Dominguez et al., 2012; Lescroart et al., 2012; Vincent and Buckingham, 2010). aSHF progenitors contribute to the right ventricular and outflow tract myocardium, while progenitor cells located in the pSHF contribute to the venous pole as well as the distal arterial pole of the heart.

A complex network of signaling inputs and transcriptional regulators is required to regulate SHF development (Rochais et al., 2009). Among these signaling molecules, retinoic acid (RA) has been shown to pattern the SHF (Stefanovic and Zaffran, 2017; Hochgreb et al., 2003). Specifically, RA signaling is required to define the posterior limit of the SHF, as indicated by the abnormal posterior expansion of the expression of aSHF markers genes, including *Fgf8*, *Fgf10*, and *Tbx1*, in *Raldh2*-mutant embryos (Ryckebusch et al., 2008; Sirbu et al., 2008). *Hoxa1*, *Hoxa3* and *Hoxb1* are expressed in overlapping sub-populations of cardiac progenitor cells in the pSHF and downregulated prior to differentiation (Bertrand et al., 2011). *Hoxb1-* and *Hoxa1-*expressing progenitor cells located in the pSHF segregate to both cardiac poles, contributing to the inflow tract and the inferior wall of the outflow tract (Lescroart and Zaffran, 2018; Bertrand et al., 2011). In contrast, cardiac progenitors that contribute to the superior wall of the outflow tract and right ventricle do not express Hox transcription factors. *Hoxb1* is required for normal deployment of SHF cells during outflow tract development (Roux et al., 2015). TALE-superclass transcription factors (three-amino acid length extension) such as Pbx1-3 or Meis1-2, which are co-factors of anterior Hox proteins, are also expressed in cardiac progenitors, suggesting a wider role for HOX/TALE complexes during SHF development (Paige et al., 2012; Wamstad et al., 2012; Stankunas et al., 2008).

Identification of SHF-restricted regulatory elements has provided evidence that different transcriptional programs operate in distinct SHF sub-populations. Cells expressing *Cre* recombinase under the control of a SHF-restricted regulatory element from the *Mef2c* gene, contribute widely to the outflow tract and right ventricle, as well as a population of cells at the venous pole of the heart giving rise to the primary atrial septum and DMP (De Bono et al., 2018; Goddeeris et al., 2008; Verzi et al., 2005; Dodou et al., 2004). Although subdomains of the SHF prefigure and are essential to establish distinct structures within the mature heart, it is unclear how distinct sub-populations are defined. Here, we identify the genome-wide transcriptional profiles and chromatin accessibility maps of sub-populations of SHF cardiac progenitor cells using RNA- and ATAC-sequencing approaches on purified cells. Through gain and loss of function experiments we identify Hoxb1 as a key upstream player in SHF patterning and deployment. Mis-expression of *Hoxb1* in the Hox-free domain of the SHF results in aberrant cellular identity of progenitor cells and arrested cardiac differentiation, leading ultimately to cell death. The addition of progenitor cells from the pSHF to the venous pole is also impaired in *Hoxa1^-/-^; Hoxb1^-/-^* hearts, resulting in abnormal development of the DMP and consequent atrioventricular septal defects (AVSDs). Hoxb1 is thus a critical determinant of cardiac progenitor cell fate in vertebrates.

## RESULTS

### Transcriptomic and epigenomic profiling of the SHF

To identify the transcriptional profiles of distinct cardiac progenitor populations, we made use of two transgenic mouse lines, *Hoxb1^GFP^* and *Mef2c-AHF-Cre* (*Mef2c-Cre*), that drive reporter gene expression in sub-domains of the SHF (Roux et al., 2015; Bertrand et al., 2011); (Briggs et al., 2013; Verzi et al., 2005). At embryonic day (E) 9.5 (16 somites [s]), the GFP reporter of *Hoxb1^GFP^* embryos is detectable in the posterior region of the SHF (Figure 1A). Genetic lineage analysis of *Hoxb1*-expressing cells using the *Hoxb1^IRES-Cre^* mouse line showed that *Hoxb1* progenitors contribute to both atria, the DMP and the myocardium at the base of the pulmonary trunk at E11.5-E12.5 (Figures 1B,C). Genetic lineage analysis of *Mef2c-Cre*-labelled cells using *Mef2c-Cre;Rosa^tdT^* mouse line showed that Tomato-positive (Tomato+) cells are detected in the arterial pole of the heart and the DMP at E9.5-E10.5 (Figures 1D,1E). At E12.5, the contribution of *Mef2-Cre*-expressing cells is observed in the great arteries (aorta and pulmonary trunk) and the right ventricle (Figure 1F), consistent with previous observations (De Bono et al., 2018; Goddeeris et al., 2008; Verzi et al., 2005). To further characterize the expression pattern of these two reporter lines we performed RNA-FISH (RNAscope fluorescent *in situ* hybridization). At E8.5-9, RNA-FISH showed that expression of *Osr1*, a gene expressed in pSHF progenitors (Zhou et al., 2015), largely overlapped with *Hoxb1* expression (Figures 1G and 1I), whereas *Mef2c-Cre* predominantly labeled a distinct progenitor cell population to *Osr1* (Figures 1G,H). Double whole-mount *in situ* hybridization identified a minor subset of cardiac progenitors co-labeled by *Hoxb1* and *Tomato* (*Mef2c-Cre;Rosa^tdT^*), likely corresponding to progenitor cells giving rise the DMP at the venous pole and inferior outflow tract wall at the arterial pole (Figures 1I-K) {Bertrand, 2011 #2;Roux, 2015 #7;Verzi, 2005 #16;Briggs, 2013 #69}.

**Figure 1:**
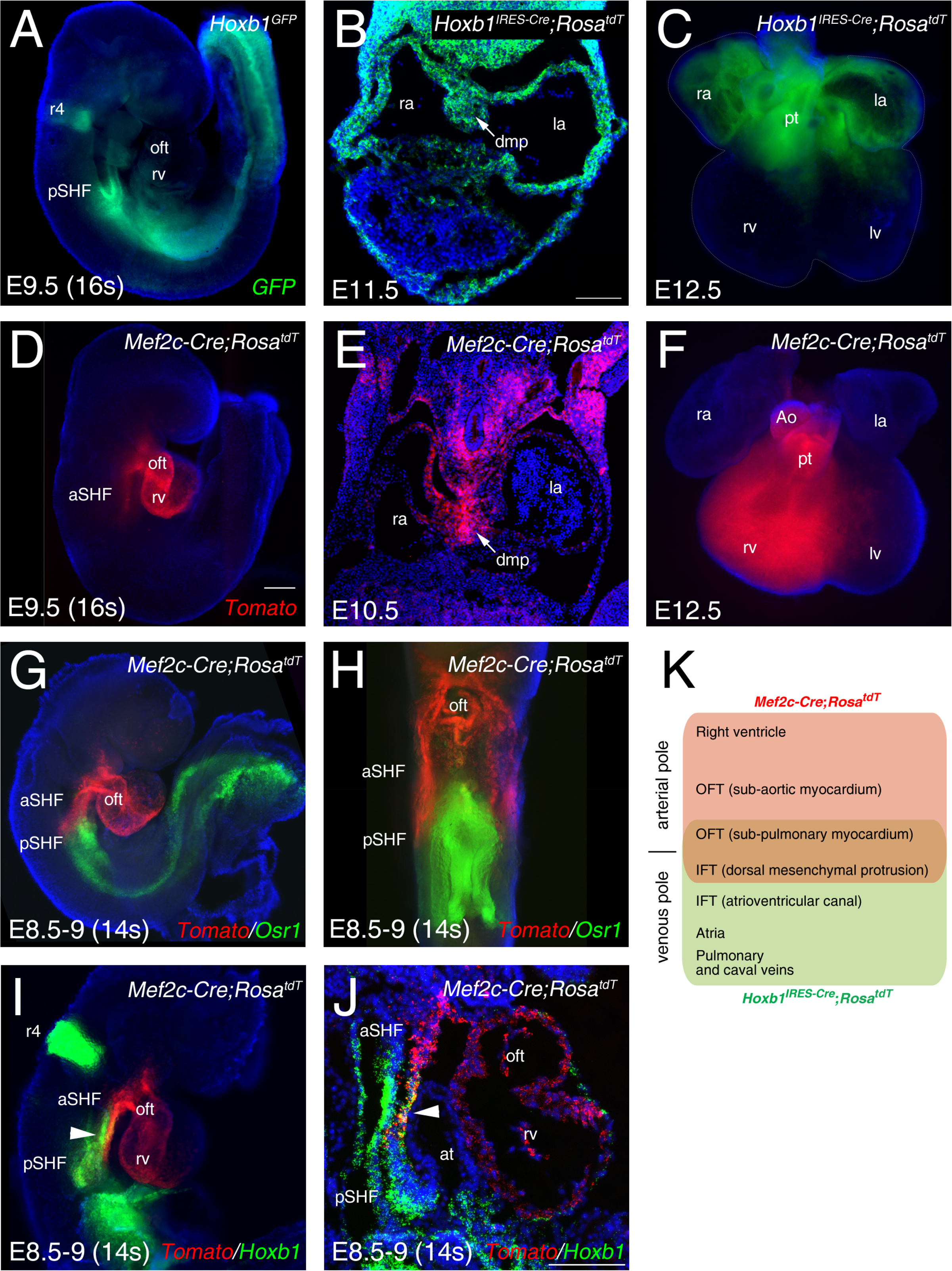
Characterization of two transgenes defining complementary domains of the SHF. (A) Whole-mount fluorescence microscopy of E9.5 (16somites [s]) *Hoxb1^GFP^* embryos. (B) Transverse section at E11.5 heart showing *Hoxb1-Cre* genetic lineage contribution to atrial myocardium and the dorsal mesenchymal protrusion (DMP). (C) Ventral view of an E12.5 heart showing the *Hoxb1-Cre* (*Hoxb1^IRES-Cre^;Rosa* - green) genetic lineage contributions to both atria and sub-pulmonary myocardium. (D) E9.5 (16s) *Mef2c-Cre;Rosa^tdT^* embryos showing the contribution of the *Mef2c-Cre* genetic lineage (Tomato, red) to the outflow tract and the right ventricle. (E) Transverse section at E10.5 showing the *Mef2c-Cre* genetic lineage contribution to the DMP. (F) Ventral view of an E12.5 heart showing the *Mef2c-Cre* genetic lineage contribution to the right ventricle and great arteries. (G) RNA-FISH showing the expression of *Osr1* (green) and the *Mef2c-Cre* labeled cells (Tomato; red). (H) Ventral view of the embryo shown in G. Distribution of *Osr1* is detected in the posterior domain of *Mef2c-Cre*. (I,J) RNA-FISH showing a small domain of overlap of *Hoxb1* (green) and *Mef2c-Cre* labeled cells (Tomato; red). (K) Cartoon summarizing the contribution of the *Hoxb1*-*Cre* (green) and *Mef2c-Cre* (red) lineages in the embryo at E9.5. Nuclei are stained with Hoechst. ao, aorta; at, atria; aSHF, anterior second heart field; avc, atrioventricular canal; ift, inflow tract; la, left atria; lv, left ventricle; oft, outflow tract; pt, pulmonary trunk; pSHF, posterior second heart field; ra, right atria; rv, right ventricle; Scale bars represent 100 μm (C, J); 200 μm (D).

After micro-dissection and dissociation of the SHF region from *Hoxb1^GFP^* and *Mef2c-Cre;Rosa^tdT^* embryos at E9.5 (16s; n=3 each), GFP+ or Tomato+ cells were purified by flow cytometry-activated cell sorting (FACS) and subsequently used for RNA-seq (Figure 2A). FACS analysis showed that Tomato+ and GFP+ cells comprise, respectively, 33% and 23% of the total micro-dissected region (Figures 2B,D). The enrichment was validated for a set of genes known to be specifically expressed in the pSHF *(Hoxa1, Hoxb1, Osr1, Tbx5* and *Aldh1a2)* (Figures 2C,E). Principal component analysis (PCA) and calculation of the Euclidean distance between the regularized log (rlog)-transformed data for each sample using DESeq2 demonstrated the strong similarity between biological replicates (Figure 2-figure supplement 1). We identified 11,970 genes expressed in both cell types. 2,133 genes were transcribed specifically in the GFP+ population (Figure 2F **and** Figure 2-figure supplement 1). Gene ontology (GO) enrichment analysis for the biological processes linked to these genes showed a significant enrichment of GO terms associated with “heart development”, “epithelium development”, “cardiac chamber morphogenesis” and “cell adhesion” (Figure 2G). Included in the “heart development” list we identified several genes previously described as being expressed in the posterior region of the SHF (*e.g.*, *Tbx5*, *Osr1*, *Tbx18*, *Foxf1,* and *Wnt2*), as well as *Bmp4*, *Nr2f2*, *Sema3c*, *Gata4* (Figure 2H- **and** Figure 2-figure supplement 2; Figure 2-supplementary file 1). RNA-FISH analysis validated the expression of *Bmp4* in the pSHF (Figure 2-figure supplement 2).

**Figure 2:**
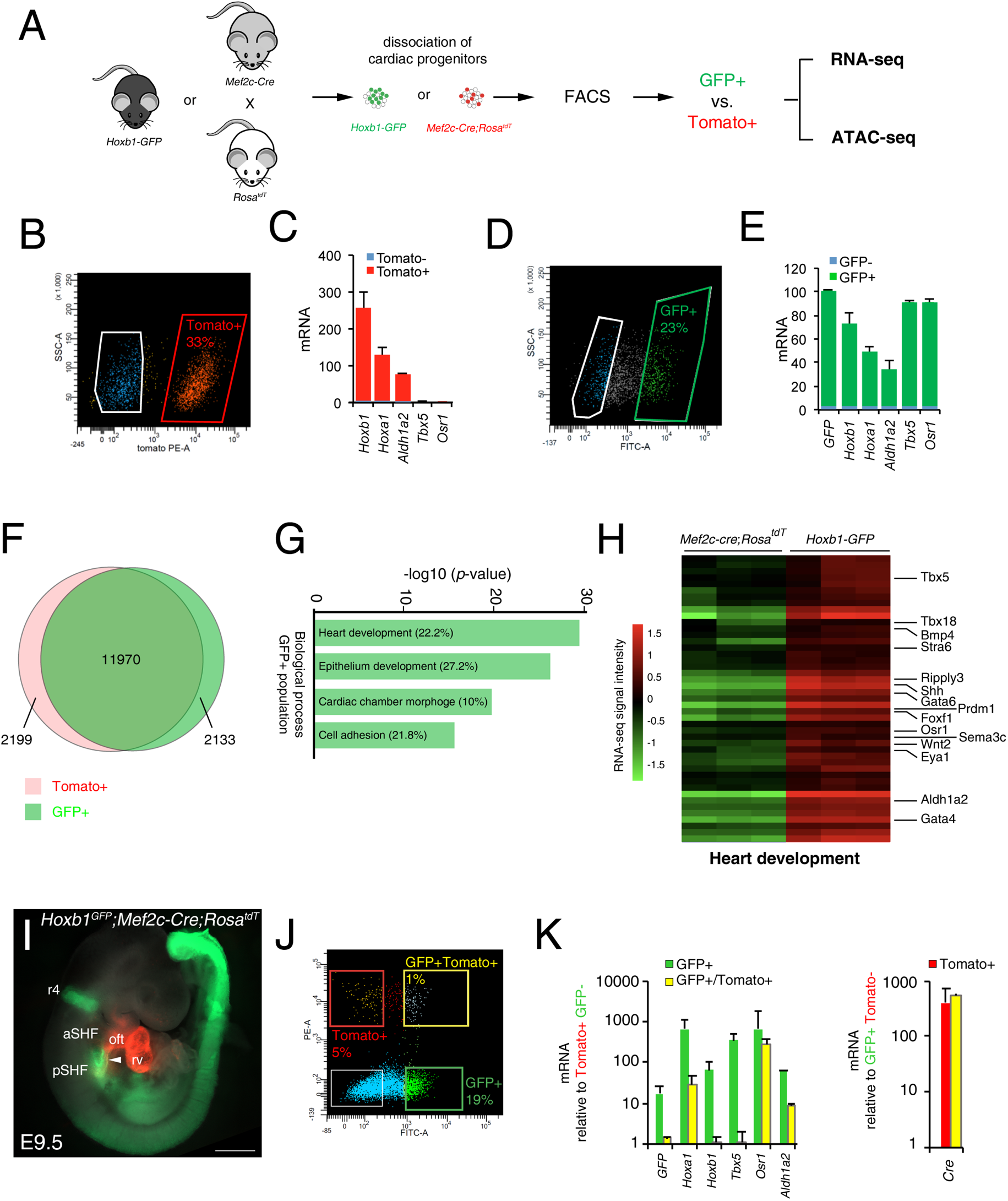
Molecular signature of the posterior SHF. (A) Scheme of the protocol utilized to characterize the molecular signature of the SHF on isolated *Mef2c-Cre;Rosa^tdT^* (Tomato) and *Hoxb1^GFP^* (GFP) positive cells. (B,D) FACS profile of E9.5 cardiac progenitor cells isolated from *Mef2c-Cre;Rosa^tdT^* and *Hoxb1^GFP^* embryos. (C,E) Expression of pSHF markers (*Hoxa1, Hoxb1, Osr1, Raldh2, Tbx5, GFP*) was analyzed with real-time qPCR. Data were normalized to *HPRT* and expressed as folds of increase over untreated samples (negative population). (F) Venn diagram showing transcripts differentially expressed in the GFP+ (green) compared to the Tomato+ (red) populations. (G) Gene ontology (GO) analysis of GFP+ progenitor cells performed with ClusterProfiler system showing enrichment of upregulated genes in the pSHF with ranked by -log_10_ (*p*-value). (H) Example of the heatmap of “heart development” GO term associated genes analyzed by RNA-seq (n=3 from GFP+ cells, n=3 from Tomato+ cells). (I) Whole-mount fluorescence microscopy of triple transgenic *Hoxb1^GFP^;Mef2c-Cre;Rosa^tdT^* embryos at stage E9.5. (J) FACS profile of E9.5 cardiac progenitor cells isolated from *Hoxb1^GFP^;Mef2c-Cre;Rosa^tdT^* embryos. (K) Expression of pSHF markers (*Hoxa1, Hoxb1, Osr1, Aldh1a2, Tbx5*), *GFP* and the *Cre* recombinase were analyzed with real-time qPCR. Data were normalized to *HPRT* and expressed as folds of increase over untreated samples (negative population). Scale bars: 500 μm.

Given the small overlap between *Hoxb1* and Tomato expression (Figure 1J), we generated triple transgenic *Hoxb1^GFP^;Mef2c-Cre;Rosa^tdT^* embryos at stage E9.5 (Figure 2I) and purified double positive (GFP+/Tomato+) and simple positive (GFP+ or Tomato+) cells. FACS analysis showed that the GFP+/Tomato+ gate comprised only 1% of the total micro-dissected region (Figure 2J). Interestingly, transcriptional analysis revealed that both *Hoxb1* and *GFP* transcripts were decreased in the GFP+/Tomato+ population compared to the GFP+ population. Conversely, *Cre* transcripts were equally expressed in both the Tomato+/GFP+ and Tomato+ populations suggesting that the *Mef2c-AHF* enhancer was still active in sub-pulmonary myocardial progenitors at this stage.

To define accessible sites for transcriptional regulation in SHF sub-populations, we performed ATAC-seq (Buenrostro et al., 2015). We performed ATAC-seq on FACS-sorted Tomato+ or GFP+ cells from E9.5 (12-14s) embryos. Samples were subjected to massively parallel sequencing and overlapping peaks from replicate samples were merged to identify high-confidence regions of open chromatin. The correlation heatmaps and PCA plot highlighted the differences in ATAC read concentrations between Tomato+ and GFP+ samples (Figure 3-figure supplement 3). ATAC-seq peaks representing open chromatin were highly reproducible between biological replicates and showed a clear enrichment at regulatory elements (Figure 3-figure supplement 3). We performed a stringent analysis to identify qualitative (present or absent peaks only) differences in chromatin accessibility. By comparing the signal for each peak in Tomato+ and GFP+ populations, we identified 1,285 peaks that were exclusively accessible in the GFP+ population (Figures 3A,B). Approximately 94% of peaks found in the GFP+ population were also present in the Tomato+ population, while 3.5% of the peaks were exclusively present in the GFP+population (Figure 3B). We then asked whether DNA regions differentially accessible between the Tomato+ and GFP+ populations were selectively associated with genes specifically expressed in the pSHF that were identified from our RNA-seq analysis. In order to assess differential chromatin accessibility at each consensus peaks we used an affinity analysis as a quantitative approach (Figure 3C). Quantitative analysis confirmed the number of peaks identified by the qualitative approach for each GFP+ or Tomato+ population. In addition, quantitative differences in peak signals were observed between the Tomato+ and GFP+ populations (Figure 3-figure supplement 3). We next asked if global differences between the accessible chromatin landscapes of the Tomato+ and GFP+ populations correlated with changes in gene expression. Thus, we mapped individual ATAC-seq peaks in the Tomato+ and GFP+ populations based on their distance to transcription start site (TSS) and examined the expression of the corresponding genes (Figures 3D,E). Changes in chromatin accessibility did not correlate precisely with changes in gene expression and several peaks near differentially expressed genes were not differentially accessible. Such decoupling between enhancer accessibility and activity has been observed in other developmental contexts, including early cardiogenesis (Racioppi et al., 2019; Paige et al., 2012; Wamstad et al., 2012). However, we found 53 (Tomato+ population) and 65 (GFP+ population) peaks correlating with changes in gene expression (Figure 3E). Among the 65 peaks specific to the GFP+ population we found *Hoxb1*, *Aldh1a2* and *Sema3c*, loci, which showed open chromatin regions concentrated in the promoter and regulatory regions occupied by several transcription factors (Figures 3D-F; 3H). ATAC-seq data for the GFP+ population thus revealed a high read count around the promoter regions of genes enriched in the pSHF, including *Hoxb1*, *Aldh1a2, Osr1* and *Sema3c* (Figures 3F-H). Similarly, the pSHF enhancer previously identified at the *Foxf1a* locus (Hoffmann et al., 2014) exhibited enrichment of ATAC-seq reads in GFP+ population (Figure 3-figure supplement 3, Figure 3-figure supplementary file 2). In contrast, the established *Mef2c* anterior heart field enhancer (*Mef2c-F6;* 285-bp) was marked by open chromatin in the Tomato+ population, but not the GFP+ population (Figure 3-figure supplement 3), confirming that ATAC-seq marks active promoters and enhancers in prominent compartment-specific patterns. Among the ATAC-seq peaks we observed a high read density within an intron of the *Mef2c* gene, approximately 2.5-kb or 4-kb upstream of the previously described *Mef2c*-AHF enhancers (Dodou et al., 2004; von Both et al., 2004).

**Figure 3:**
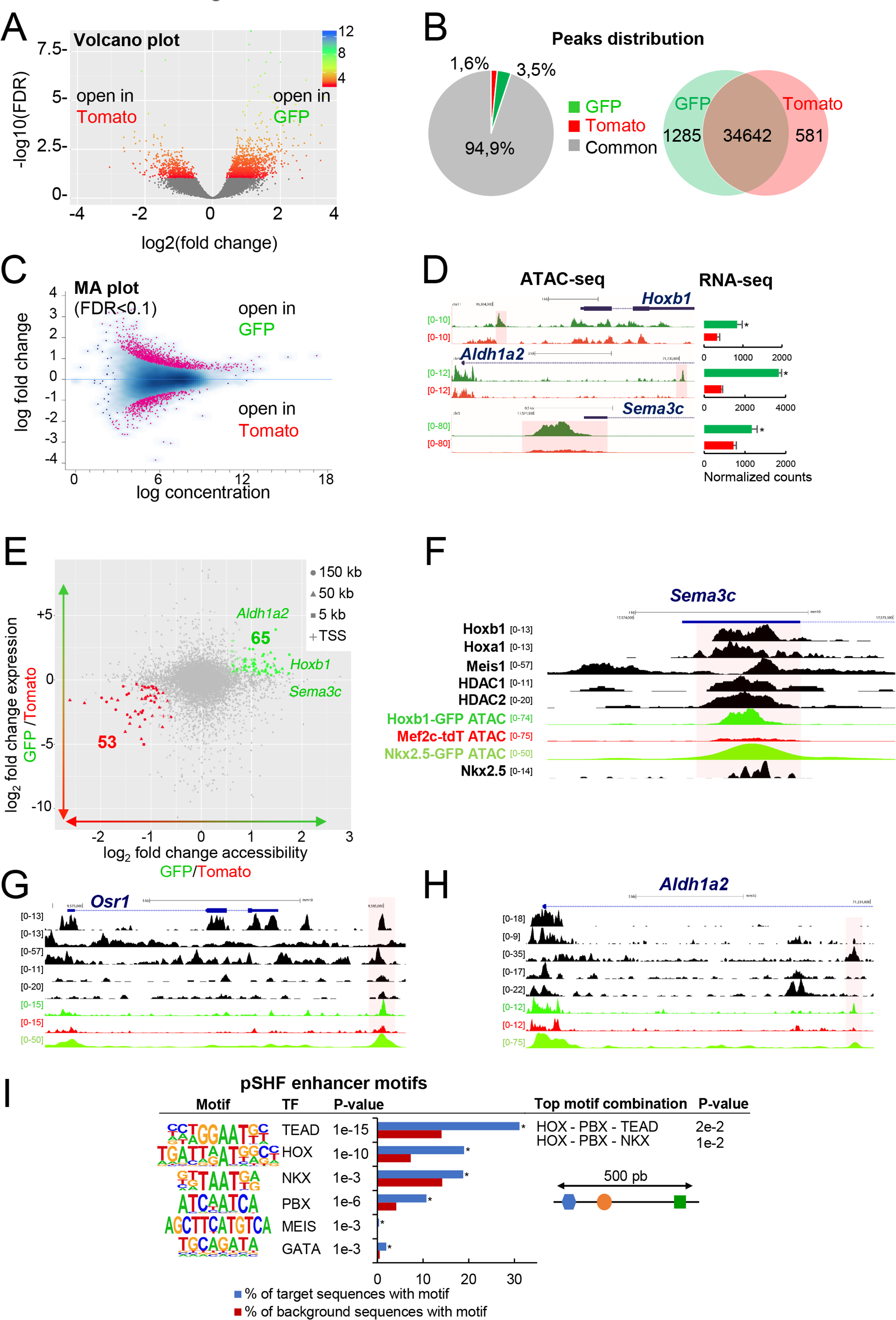
Differential chromatin accessibility in GFP+ and Tomato+ cardiac progenitor cells. (A) Volcano plot of ATAC-seq performed from GFP+ and Tomato+ cardiac progenitors. (B) Pie chart showing the distribution of the ATAC-seq peaks in the two populations. (C) MA plot of ATAC-seq peaks in GFP+ *versus* Tomato+ cells. (D) Open chromatin profiles correlate with transcriptional expression. Browser views of *Hoxb1*, *Aldh1a2* and *Sema3c* loci with ATAC-seq profiles of GFP+ pSHF progenitor cells (green) and Tomato+ *Mef2c-Cre* labeled cells (red). Data represent merged technical and biological replicates for each cell type. The *y*-axis scales range from 0–80 in normalized arbitrary units. The tracks represent ATAC-seq, whereas the bar graphs represent RNA-seq. Boxed regions show cell-type-specific peaks around *Aldh1a2, Osr1, and Sema3C* gene loci. (E) Change in accessibility versus change in gene expression in GFP+ and Tomato+ cells. For each ATAC-seq peak, the log of the ratio of normalized ATAC-seq reads (GFP/Tomato) is plotted on the x-axis, and the log of the ratio normalized RNA-seq reads corresponding to the nearest gene is plotted in the y-axis. Peaks that are both significantly differentially accessible (FDR<0.1) and significantly differentially expressed (FDR<0.1) are colored green (more open in GFP+ cells, higher expression in GFP+ cells; 65 peaks) or red (more open in Tomato population, higher expression in Tomato; 53 peaks). (F-H) Browser views of *Sema3c* (F), *Osr1* (G) and *Aldh1a2* (H) gene loci with ATAC-seq profiles of GFP+ pSHF progenitor cells (green) and Tomato+ *Mef2c-Cre* labeled cells (red). Open chromatin profiles correlate with transcription factor binding at putative enhancers specific for cardiac progenitor cells (I) pSHF enhancers are enriched in DNA binding motifs for HOX and known cardiac transcription factors. HOX recognition motifs were strongly enriched in a known motif search in pSHF enhancers. Other enriched matrices match binding sites of known cardiac regulators. HOX binding motifs are highly enriched at genomic regions bound by cardiac transcription factors. *p*-values were obtained using HOMER (Heinz et al., 2010). Combinations of 3 sequence motifs contained within 500-bp are shown.

Together this analysis indicates that our dataset can be used to identify regulatory elements in distinct SHF sub-populations. ATAC-seq in GFP+ cells identified several pSHF-specific peaks indicating that these regions may function as enhancers (Figures 3F-H). The most highly enriched motif in ATAC-seq regions of open chromatin was the consensus Hox motif (Figure 3I) (Fan et al., 2012). Other significantly enriched motifs include putative binding sites for Pbx and Meis proteins (Figure 3I), TALE-class transcription factors interacting with Hox proteins (Lescroart and Zaffran, 2018; Ladam and Sagerstrom, 2014). Members of the Pbx and Meis families have been previously identified as cofactors of Hoxb1 in mammalian cell lines and embryonic tissues, indicating a high level of functional conservation (Roux and Zaffran, 2016; Ladam and Sagerstrom, 2014; Mann et al., 2009). In addition, we found overrepresentation of motifs for GATA and TEA domain (TEAD) family transcription factors as well as for the Nkx2-5 homeodomain. Because transcription factors function in a combinatorial manner, we identified combinations of multiple motifs that were most enriched at pSHF candidate enhancers relative to non-pSHF enhancers (Figure 3I). Our computational analysis showed that the most enriched combinations contained Hox motifs adjacent to TALE transcription factor recognition sequences. Consistent with these observations, chromatin immunoprecipitation (ChIP)-sequence data for the cofactors of Hox proteins (Meis1, Nkx2-5, HDACs) revealed that these factors bind putative regulatory elements marked by open chromatin in the GFP+ but not in the Tomato+ population (Figures 3F-H).

### Mis-expression of *Hoxb1* in the *Mef2c-AHF-Cre* lineage disrupts right ventricular formation

In order to investigate the role of Hoxb1 during heart development we generated a conditional activated *Tg*(*CAG-Hoxb1-EGFP)^1Sza^* (*Hoxb1^GoF^*) transgenic mouse, in which *Hoxb1* cDNA expression is activated upon Cre-mediated recombination (Zaffran et al., 2018). *Hoxb1^GoF^* mice were crossed with to *Mef2c-AHF-Cre* (*Mef2c-Cre*) mice to mis-express *Hoxb1* in *Mef2c-AHF+* cells (Verzi et al., 2005). *Hoxb1^GoF^;Mef2c-Cre* embryos exhibited severe heart defects as early as E9.5, as observed by a looping defect and common ventricular chamber in transgenic compared to control embryos (Figures 4A-B). Expression of GFP in the aSHF and its derivatives confirmed *Mef2c-Cre-*driven recombination (Figures 4A’,B’). Immunostaining of E10.5 control embryos revealed normal future right and left ventricular chambers with developing trabeculae (Figure 4C); in *Hoxb1^GoF^;Mef2c-Cre* embryos, the heart was abnormally shaped with no clear distinction between right and left ventricular chambers (Figure 4D). The phenotype was even more pronounced at E12.5, when H&E-stained transverse sections showed a hypoplastic right ventricle with abnormal positioning of the ventricular septum (Figures 4E, 4F). Hypoplasia of the right ventricle in *Hoxb1^GoF^;Mef2c-Cre* embryos was evident at E15.5 (Figures 4G-J; 8/8). At this stage, an abnormally thin right ventricular wall was observed (5/8). In addition, 50% of *Hoxb1^GoF^;Mef2c-Cre* embryos showed misalignment of the great arteries (4/8) and 63% displayed ventricular septal defects (VSD; 5/8). Overall, these results suggest that ectopic *Hoxb1* expression in the *Mef2c-AHF* lineage disrupts the contribution of anterior cardiac progenitor cells to the forming heart.

**Figure 4:**
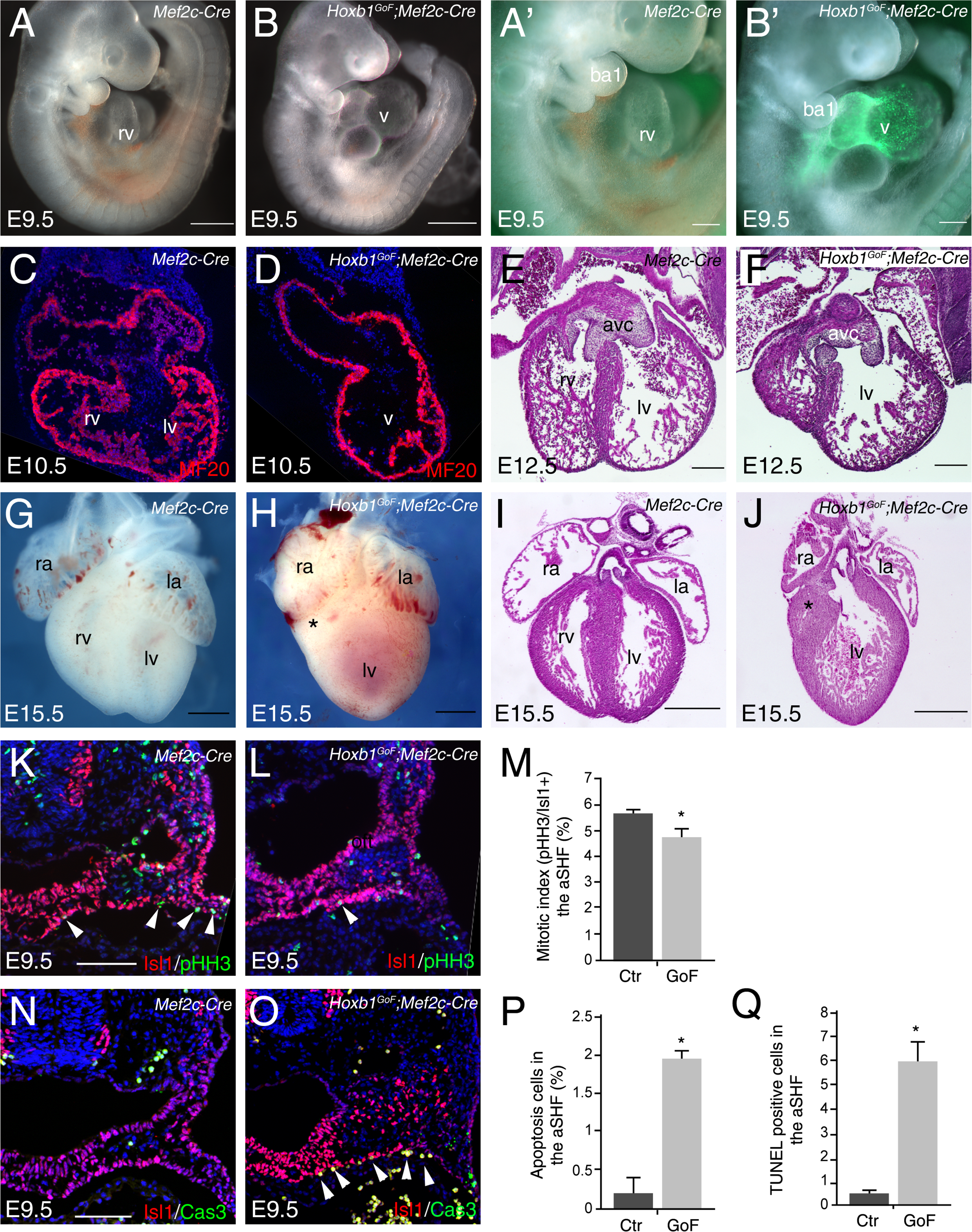
Activation of *Hoxb1* expression in aSHF progenitors disrupts the formation of the right ventricle. (A,B) Macroscopic view of *Mef2c-AHF-Cre* (*Mef2c-Cre*, control) and *Hoxb1^GoF^*;*Mef2c-Cre* embryos at E9.5. (A’,B’) High magnification of embryo in a and b showing GFP activity in the Cre-recombinase driven cells. (C,D) Immunofluorescence with MF20 (red) in control (C) and *Hoxb1^GoF^*;*Mef2c-Cre* (D) hearts at E10.5. (E-J) Hematoxylin & Eosin (H&E) staining on transversal section of control (E,I) and *Hoxb1^GoF^*;*Mef2c-Cre* (F,J) hearts at E12.5 and E15.5. The right ventricle (asterisk) in *Hoxb1^GoF^*;*Mef2c-Cre* hearts is hypoplastic compared to control hearts (n=8). (G,H) Whole-mount views of control (G) and *Hoxb1^GoF^*;*Mef2c-Cre* (H) hearts at E15.5. (K,L) Immunofluorescence with Isl1 (red) and Phospho-histone H3 (pHH3, green) on *Mef2c-Cre* (K) and *Hoxb1^GoF^*;*Mef2c-Cre* (L) embryos at E9.5. (M) Quantification of pHH3-positive cells in the aSHF Isl1+, showed a reduced of the mitotic index in *Hoxb1^GoF^*;*Mef2c-Cre* (n=3) compared to *Mef2c-Cre* (*Ctr*) (n=7) embryos. (N,O) Immunofluorescence with Isl1 (red) and Caspase 3 (Cas3, green) on *Mef2c-Cre* (N) and *Hoxb1^GoF^*;*Mef2c-Cre* (O) embryos at E9.5. Arrowheads indicate Cas3-positive cells. (P) Quantification of Cas3-positive cells revealed increased cells death in the aSHF of *Hoxb1^GoF^*;*Mef2c-Cre* embryos. (Q) Quantification of TUNEL staining performed on *Mef2c-Cre* (*Ctr*) and *Hoxb1^GoF^*;*Mef2c-Cre* embryos. ao, aorta; avc, atrioventricular canal; ba, branchial arch; la, left atrium; lv, left ventricle; oft, outflow tract; PM, pharyngeal mesoderm; pt, pulmonary trunk; ra, right atrium; rv, right ventricle. Scale bars: 100μm(A’,B’,E,F); 200μm(A’,B’,E,F); 500μm(A,B,G-J).

### Ectopic *Hoxb1* activity affects the survival of SHF progenitor cells

To study the cause of the right ventricular hypoplasia defect in *Hoxb1^GoF^;Mef2c-Cre* embryos, we performed a lineage analysis of aSHF progenitors using the *Rosa26R-lacZ* (*R26R*) reporter line. By E9, β-galactosidase (β-gal)-positive cells were detected in the SHF, the outflow tract and right ventricle of control embryos, reflecting the normal contribution of cardiac progenitor cells (Figure 4-figure supplement 4). In contrast, a striking reduction of the β-gal+ expression domain was observed in *Hoxb1^GoF^;Mef2c-Cre;R26R* embryos (Figure 4-figure supplement 4). The reduction was evident within the heart and in the SHF of these embryos (Figure 4-figure supplement 4; asterisk). This observation was further confirmed using the *Mlc1v-nlacZ-24* (*Mlc1v24*) transgenic line, containing an *Fgf10* enhancer trap transgene expressed in the aSHF (Figure 4-figure supplement 4) Quantitative analysis demonstrated a reduction of the β-gal+ expression domain in the *Hoxb1^GoF^;Mef2c-Cre;Mlc1v24* compared to control embryos consistent with a reduction of the distance between the arterial and venous poles (Figure 4-figure supplement 4). These data reveal that cardiac defects in *Hoxb1^GoF^;Mef2c-Cre* embryos are associated with a decrease in progenitor cell numbers in the aSHF.

To determine the origin of this decrease we performed proliferation and cell death assays at E9.5. More specifically, we determined the mitotic index (pHH3+ cells) of Isl1+ cardiac progenitor cells. There was a modest reduction in cell proliferation between control and *Hoxb1^GoF^;Mef2c-Cre* embryos (Figures 4K-M). However, a significant increase of Caspase-3 and TUNEL-positive cells was detected in the aSHF (Isl1+) of *Hoxb1^GoF^;Mef2c-Cre* compared to control embryos, indicating increased apoptosis (Figures 4N-Q). Hence, the severe right ventricular hypoplasia resulting from ectopic *Hoxb1* expression was primarily due to extensive cell death and secondarily to reduced proliferation in the aSHF.

### Mis-expression of *Hoxb1* in the *Mef2c-AHF+* domain disturbs cardiac progenitor identity and blocks cardiac differentiation

To better understand the change of the transcriptional program in the aSHF upon ectopic expression of *Hoxb1*, we performed RNA-seq analysis on dissected progenitor regions from control and *Hoxb1^GoF^;Mef2c-Cre* embryos (n=3 each) at E9.5 (16-20s) (Figure 5-figure supplement 5). We identified 1,378 genes upregulated and 1,345 genes downregulated in the SHF of *Hoxb1^GoF^;Mef2c-Cre* embryos. GO enrichment analysis for the biological processes associated with the upregulated genes showed significant enrichment of the GO terms “cell death” and “apoptotic signaling pathway” (Figure 5B and Figure 5-figure supplement 5), consistent with the decrease of aSHF cell numbers and increase of Cas3+ or TUNEL+ cells observed in *Hoxb1^GoF^;Mef2c-Cre* embryos (Figures 4N-Q). These genes included regulators of programmed cell death (*e.g., Bad*, *Bmf*, *Trp53,* and *Dapk3;* n=214) as well as modulators of growth and RA signaling (*e.g., Gsk3a, Crabp2, and Rarα*) (Figure 5-figure supplement 5). GO analysis of the downregulated genes revealed an enrichment in the GO terms “heart development” and “muscle cell differentiation” suggesting an inhibition of cardiac differentiation (Figure 5B and Figure 5-figure supplement 5). *Fgf10*, a well-characterized marker of the aSHF (Kelly et al., 2001), was among the most significant downregulated genes in *Hoxb1^GoF^;Mef2c-Cre* embryos (*p*=0.002) (Figure 5A and Figure 5-figure supplement 5; Figure 5A-supplementary file 1). This finding is consistent with the decrease in *Mlc1v-nlac-24* transgene expression (Figure 4-figure supplement 4) and reduction of arterial and venous pole distance measured in *Hoxb1^GoF^;Mef2c-Cre* embryos (Figure 4-figure supplement 4). Among the upregulated genes, we found *Osr1* and *Tbx5,* known to regulate cell cycle progression in the pSHF (Figures 5C and figure supplement 5; Figure 5-file supplement 1) (Zhou et al., 2015). The upregulation of these genes was confirmed by qPCR and *in situ* hybridization (Figures 5D-K and Figure 5-figure supplement 5). Analysis of *Osr1* and *Tbx5* expression profiles showed an anterior shift of their expression in *Hoxb1^GoF^;Mef2c-Cre* compared to control embryos (Figure 5D-K). Moreover, we found that the pSHF markers *Tbx5* and *Osr1* are both expressed in the *GFP+* cells (*Hoxb1^GoF^;Mef2c-Cre*) indicating that the ectopic expression of *Hoxb1* in the *Mef2c-AHF*+ domain alters cardiac progenitor cell identity (Figure 5E,E’-K,K’’).

**Figure 5:**
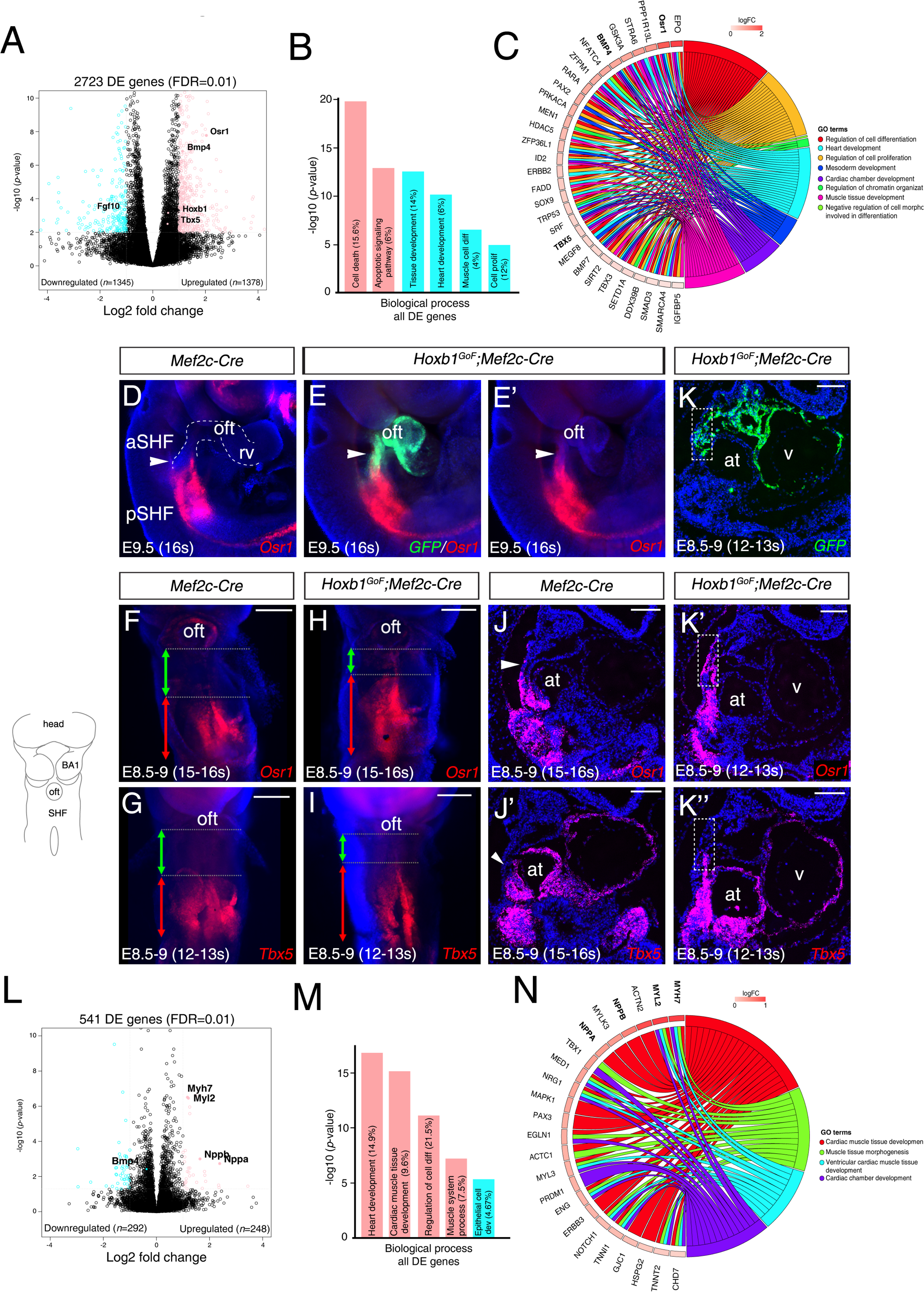
Hoxb1 regulates progenitor identity and differentiation in the pSHF. (A) Volcano plot of transcriptional profiling results with significantly dysregulated genes between *Hoxb1^GoF^;Mef2c-Cre* and control SHF. The y-axis corresponds to the mean expression value of log_10_ (*p*-value), and the x-axis displays the log2 fold change value. The colored dots represent the significantly differential expressed transcripts (*p*<0.05); the black dots represent the transcripts whose expression levels did not reach statistical significance (*p*>0.05). Differential expression analysis performed using DESeq2 revealed 2,723 genes with Log_2_-fold changes ≥1 at a False Discovery Rate (FDR) ≤0.01. (B) Gene ontology (GO) analysis of genes deregulated in *Hoxb1^GoF^;Mef2c-Cre* embryos performed with ClusterProfiler system. (C) Chord plot showing a selection of genes upregulated in *Hoxb1^GoF^;Mef2c-Cre* embryos present in the represented enriched GO terms. Outer ring shows log2 fold change or GO term grouping (right, key below). Chords connect gene names with GO term groups. (D) Whole-mount RNA-FISH for *Osr1* on E9.5 *Mef2c-Cre* embryos in lateral views. (E, E’) Whole-mount RNA-FISH for *GFP* (green) *and Osr1* (red) on E9.5 *Hoxb1^GoF^;Mef2c-Cre* embryos. Whole-mount RNA-FISH for *Osr1* in ventral views (F, H) and *Tbx5* (G, I) on E8.5-9 *Mef2c-Cre* and *Hoxb1^GoF^;Mef2c-Cre* embryos. An anteriorly shifted expression of *Osr1* and *Tbx5* is detected in *Hoxb1^GoF^;Mef2c-Cre* embryos compared to their control littermates (same somite stage – 15-16s for *Osr1* and 12-13s for *Tbx5*). RNA-FISH against *Osr1*, *Tbx5* and *GFP* on serial sections in *Hoxb1^GoF^;Mef2c-Cre* embryos (K, K’, K’’) compared to control littermates (J, J’; Serial sagittal sections). (L) Volcano plot showing differential expressed genes between *Hoxb1^-/-^* and wild-type samples. The y-axis corresponds to the mean expression value of log_10_ (*p*-value), and the x-axis displays the log2 fold change value. Colored dots represent the significantly differential expressed transcripts (*p*<0.05); the black dots represent the transcripts whose expression levels did not reach statistical significance (*p*>0.05). We identified 249 genes upregulated, and 292 genes downregulated in *Hoxb1^-/-^* embryos. (M) GO analysis of genes deregulated in *Hoxb1^-/-^* embryos with ranked by -log_10_ (*p*-value). (N) Chord plot showing a selection of genes upregulated in dissected pharyngeal mesoderm of *Hoxb1^-/-^* embryos present in the represented enriched GO terms. Outer ring shows log2 fold change or GO term grouping (right, key below). Chords connect gene names with GO term groups. Nuclei are stained with Hoechst. BA1, branchial arch 1; oft, outflow tract; SHF, second heart field. Scale bars: 200 μm (F, G, H, I); 100 μm (J, J’, K, K’, K’’).

### Hoxb1 loss of function leads to premature differentiation in the SHF

To complement our functional analysis, we determined if the cardiac differentiation program was affected in the absence of *Hoxb1* function. RNA-seq transcriptional profiling was performed on progenitor regions isolated from E8.5 (6-8s) wild-type and *Hoxb1*-deficient embryos (n=2 each; Figure 5-figure supplement 6). Interestingly, GO term analysis of the upregulated genes revealed a significant enrichment of terms including “heart development”, “cardiac muscle tissue development” and “regulation of cell differentiation” (Figures 5M and Figure 5-figure supplement 6). Upregulation of several myocardium-specific genes (*e.g., Myl2, Myh7, Actn2, Myl3, Nppa (Anf), and Nppb (Bnf)*), indicates a premature cardiac differentiation in the SHF of *Hoxb1^-/-^* mutant embryos (Figure 5N). The upregulation of these myocardial genes was confirmed by qPCR (Figure 5-figure supplement 6). The GO term “epithelial cell development” was significantly enriched in the downregulated genes (e.g., *Cdh1*, *Llgl2, and Lrp5*) (Figures 5M and Figure 5-figure supplement 6), consistent with both the deregulation of the epithelial properties of SHF cells (Cortes et al., 2018) and premature differentiation of cardiac progenitor cells (Soh et al., 2016). This loss-of-function analysis complements and supports the conclusions of our gain-of-function analysis and identifies a role for Hoxb1 in delaying differentiation and regulating progenitor cell identity in the SHF.

### Abnormal development of the SHF results in AVSD in *Hoxa1^-/-^;Hoxb1^-/-^* embryos

The formation of a transcriptional boundary between arterial and venous pole progenitor cells in the SHF has recently been shown to reflect the dynamic expression of two genes encoding T-box transcription factors, *Tbx1* and *Tbx5* (De Bono et al., 2018). Immunofluorescence analysis of E9.5 (22-23s) embryos confirmed the complementary expression of Tbx1 and Tbx5 proteins in the SHF (Figure 6A). Tbx5 expression is restricted to cells in the pSHF close to the inflow tract, whereas Tbx1 is detected in SHF cells close to the outflow tract. In *Hoxb1^GoF^;Mef2c-Cre* embryos the relative length of the Tbx1+ region close to the outflow tract revealed a significant reduction in the size of the Tbx1+ versus Tbx5+ domains (Figures 6A,B), although the boundary between Tbx1 and Tbx5-domains was established. We further analyzed the expression of Tbx1 and Tbx5 in *Hoxb1*-mutant embryos. The Tbx5+ domain appears slightly shorter in *Hoxb1^-/-^* embryos than in *Hoxb1^+/-^* littermates (Figures 6C,D). Due to redundancy between *Hoxa1* and *Hoxb1* we performed RNA-FISH analysis in compound *Hoxa1^-/-^;Hoxb1^-/-^* embryos. We found that *Tbx5*+ domain was shorter in double *Hoxa1^-^ ;Hoxb1^-/-^* compared to *Hoxb1^+/-^* littermate embryos (Figures 6E,F). These experiments suggest that forced activation of *Hoxb1* in the *Mef2c-AHF+* domain perturbs development of the aSHF. We subsequently investigated posterior SHF contributions to the venous pole of *Hoxa1^-/-^;Hoxb1^-/-^* hearts. Characterization of cardiac morphology in *Hoxa1^-/-^;Hoxb1^-/-^* hearts at fetal stages revealed lack of the DMP, a posterior SHF derivative, resulting in a primum type atrial septal defect, a class of AVSD (3/3; Figures 6G,H), Inappropriate differentiation of SHF cells may contribute to the loss of DMP formation in these mutants, providing the first evidence that *Hoxa1* and *Hoxb1* are required for atrioventricular septation.

**Figure 6:**
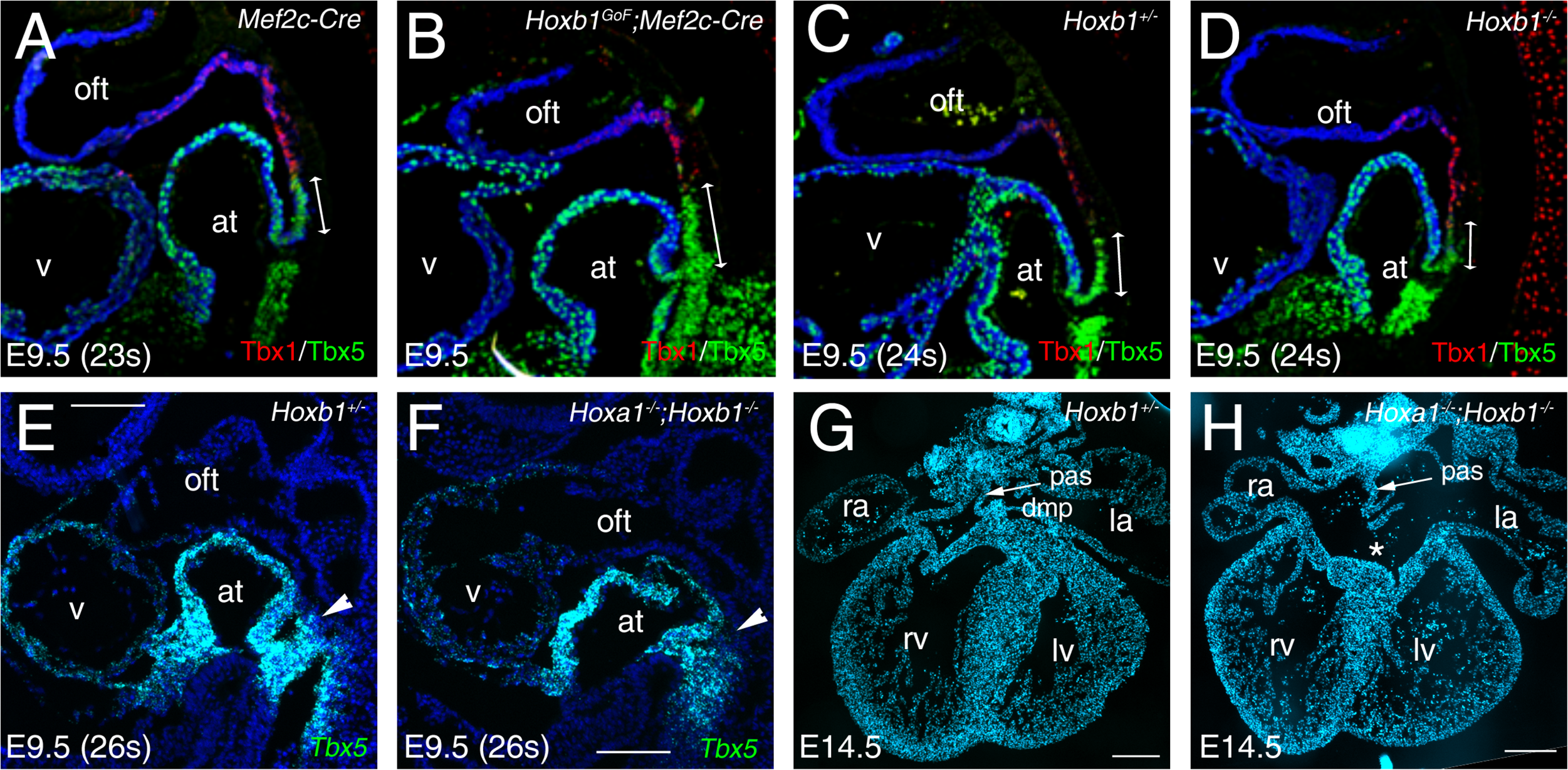
Hox genes are required for atrioventricular septation. (A-D) Immunofluorescence on medial sagittal sections showing Tbx1 (red) and Tbx5 (green) protein distribution at E9.5 (23-24s). (A,B) At E9.5, a boundary is observed in *Mef2c-Cre* embryos between Tbx1+ cells close to the arterial pole of the heart and Tbx5+ cells in the pSHF. In *Hoxb1^GoF^;Mef2c-Cre* embryos the Tbx1+ domain appears reduced although the boundary is maintained. (C,D) The Tbx5+ domain appears reduced in *Hoxb1-/-* embryos compared to *Hoxb1^+/-^* littermates. (E, F) RNA-FISH on sagittal sections showing the reduction of *Tbx5*+ domain in the pSHF of *Hoxa1^-/-^;Hoxb1^-/-^* embryos compared to *Hoxb1^+/-^* littermates. at, atria; la, left atrium; lv, left ventricle; ra, right atrium; rv, right ventricle; SHF, second heart field; v, ventricle. DAPI stained sections of a *Hoxb1+/-* (G) and a *Hoxa1-/- ;Hoxb1-/-* (H) heart at E14.5 showing the primary atrial septum (pas, arrow). Note the AVSD and absence of the DMP in H, n=3. la, left atrium; lv, left ventricle; ra, right atrium; rv, right ventricle. Scale bars: 100μm (E,F); 200μm (G,H).

### Hoxb1 is a key regulator of cardiac differentiation

We next sought to investigate the function of Hoxb1 upon cardiac induction of mouse embryonic stem (mES) cells (Figure 7A). Using a time-course gene expression analysis during cardiac differentiation of mES cells, we detected a peak of *Hoxb1* expression at day 4-5 just before the onset of cardiac differentiation (Figure 7B), as determined by the activation of specific cardiac markers such as *Myh6*, *Myh7*, *Mlc2a* and *Mlc2v* (Figure 7-figure supplement 7). Consistent with the initial activation of *Hoxb1* and *Hoxa1* during heart development in the mouse (Bertrand et al., 2011), the peak of *Hoxa1* expression was detected after day 5 (Figure 7B).

**Figure 7:**
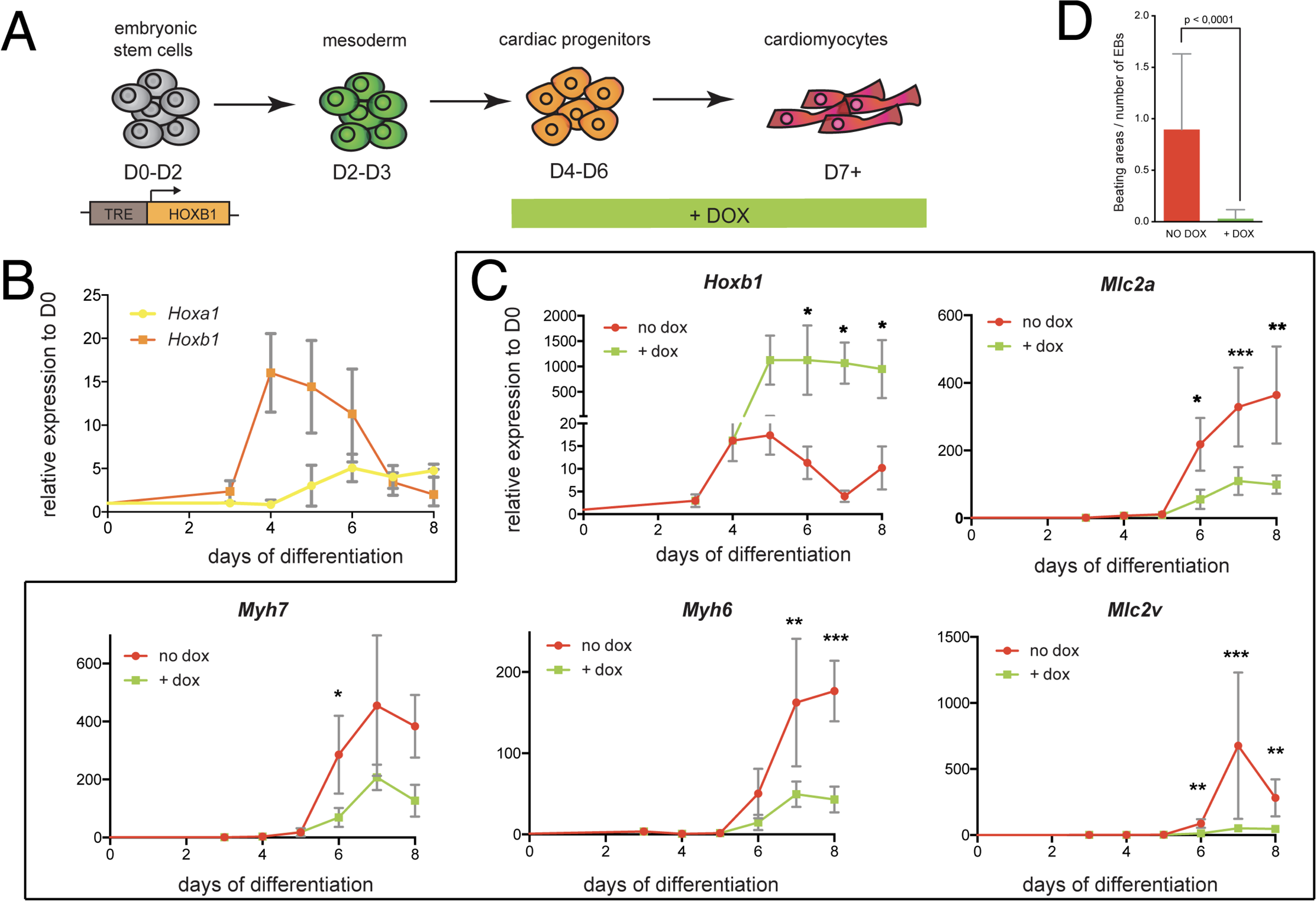
Hoxb1 overexpression in mES leads to arrested cardiac differentiation. (A) Scheme of the experiment. Using the *tet-ON/Hoxb1* mouse embryonic stem cell line (Gouti and Gavalas, 2008), *Hoxb1* expression was induced by addition of 1μg/ml of doxycycline (DOX) at the cardiac progenitor stage from day 4 (D4) during mES cell differentiation into cardiac cells. (B) Kinetics of *Hoxa1* and *Hoxb1* during mES cell differentiation as measured by RT-quantitative PCR. Results are normalized for gene expression in undifferentiated mES cells (D0). (C) Kinetics of expression for *Hoxb1* and *Myh6*, *Myh7*, *Mlc2v (or Myl2)*, *Mlc2a* during mES cell differentiation after induction of *Hoxb1* expression (+ dox) or in control condition (no dox). Results are normalized for gene expression in undifferentiated mES cells (D0). Paired, one-sided t-test was performed based on relative transcript expression between control (no dox) and doxycycline treatment (+ dox). * indicates a significance level of p<0.05, ** indicates p<0.005, *** indicates p<0.0005. (D) Quantification of beating areas relative to the number of embryonic bodies (EBs) at D8 of mES cell differentiation with or without doxycycline addition. Error bars indicate mean +/- SEM; n=4 experiments.

Next, we challenged the system by inducing continuous *Hoxb1* overexpression using the mES^Tet-on/Hoxb1^ line (Gouti and Gavalas, 2008). Permanent DOX (1 or 0.2μg/ml) treatment from day 4 of direct cardiac induction onwards (Figure 7C **and** Figure 7-figure supplement 8) interfered with the differentiation process of cardiac cells as shown by specific downregulation of the expression of *Myh7*, *Myh6*, *Mlc2a,* and *Mlc2v* (Figure 7C **and** Figure 7-figure supplement 8). Accordingly, in these conditions, we found a decreased number of beating embryoid bodies (EBs) (Figure 7D-figure supplement 8). Absence of upregulation of cell death markers attested that that reduction of beating EBs was not caused by ES cell apoptosis (Figure 7D **and** Figure 7-figure supplement 7). Interestingly, we also observed an upregulation of *Osr1*, *Tbx5* and *Bmp4* expression under these conditions, suggesting that the cellular identity of differentiating EB cells is changed, consistent with our *in vivo* observations (Figures 5C-K and Figure 7-figure supplement 7). Therefore, our data suggest that *Hoxb1* activation in mES cells results in arrest of cardiac differentiation and failure of proper identity commitment, consistent with *in vivo* results.

### Hoxb1 represses the expression of the differentiation marker *Nppa*

To identify how Hoxb1 controls cardiac differentiation, we analyzed the regulation of *Nppa* and *Nppb* expression, two markers of chamber-specific cardiomyocytes (Houweling et al., 2005). RNA-seq data showed a higher read count for *Nppa* and *Nppb* in the *Hoxb1^-/-^* compared to control embryos (Figure 8A). At E9.5, RNA-FISH confirmed an ectopic expression of *Nppa* in the SHF of *Hoxb1^-/-^* (Figures 8B,C) and *Hoxa1^-/-^Hoxb1^-/-^* (Figures 8D,E) embryos. In contrast, upon *Hoxb1* induction, the expression of *Nppa* and *Nppb* was downregulated in EBs (Figure 8-figure supplement 9). Therefore, we hypothesized that *Nppa* and *Nppb* may negatively regulate by Hoxb1 in the pSHF. ChIP-seq data in mouse ES cell lines had shown that Hoxa1, a paralog of Hoxb1, and HDAC-1 and −2 bind the *Nppa* and *Nppb* loci (De Kumar et al., 2017; Whyte et al., 2012) (Figure 8F). Coherent with these observations, we found potential Hox binding sites in a 0.7-kb *Nppa* fragment previously shown to be responsible for the expression of the *Nppa* in the developing heart (Habets et al., 2002). Thus, we hypothesized that *Nppa* and *Nppb* may be direct Hoxb1 target genes in the pSHF. As described (Durocher et al., 1997), transfection of Nkx2-5 alone or co-transfection of Nkx2-5 and Gata4 resulted in strong activation of the 0.7-kb *Nppa* promoter containing an Hox-motif in both Cos-7 and NIH3T3 cells (Figures 8G and Figure 8-figure supplement 9). However, this activity decreased threefold upon co-transfection with a Hoxb1 expression vector or co-transfection of Hoxa1 and Hoxb1, which are co-expressed in pSHF progenitor cells *in vivo* (Figure 8G), demonstrating the repressive role of Hoxb1 on the 0.7-kb *Nppa* promoter. We next assessed the activity of the 0.7-kb *Nppa* promoter in cells treated with trichostatin-A (TSA) an inhibitor of the Hdac activity known to regulate HOX functions (McKinsey, 2012). Hdacs inhibition increased the luciferase activity of the reporter constructs (Figure 8H). When co-transfected, Hoxa1 or Hoxb1 suppressed the TSA-mediated activation of the *Nppa* promoter in cell culture (Figure 8H). Together these results suggest that Hoxb1 inhibits pSHF cell differentiation by directly repressing myocardial gene transcription even under conditions of histone acetylation.

**Figure 8:**
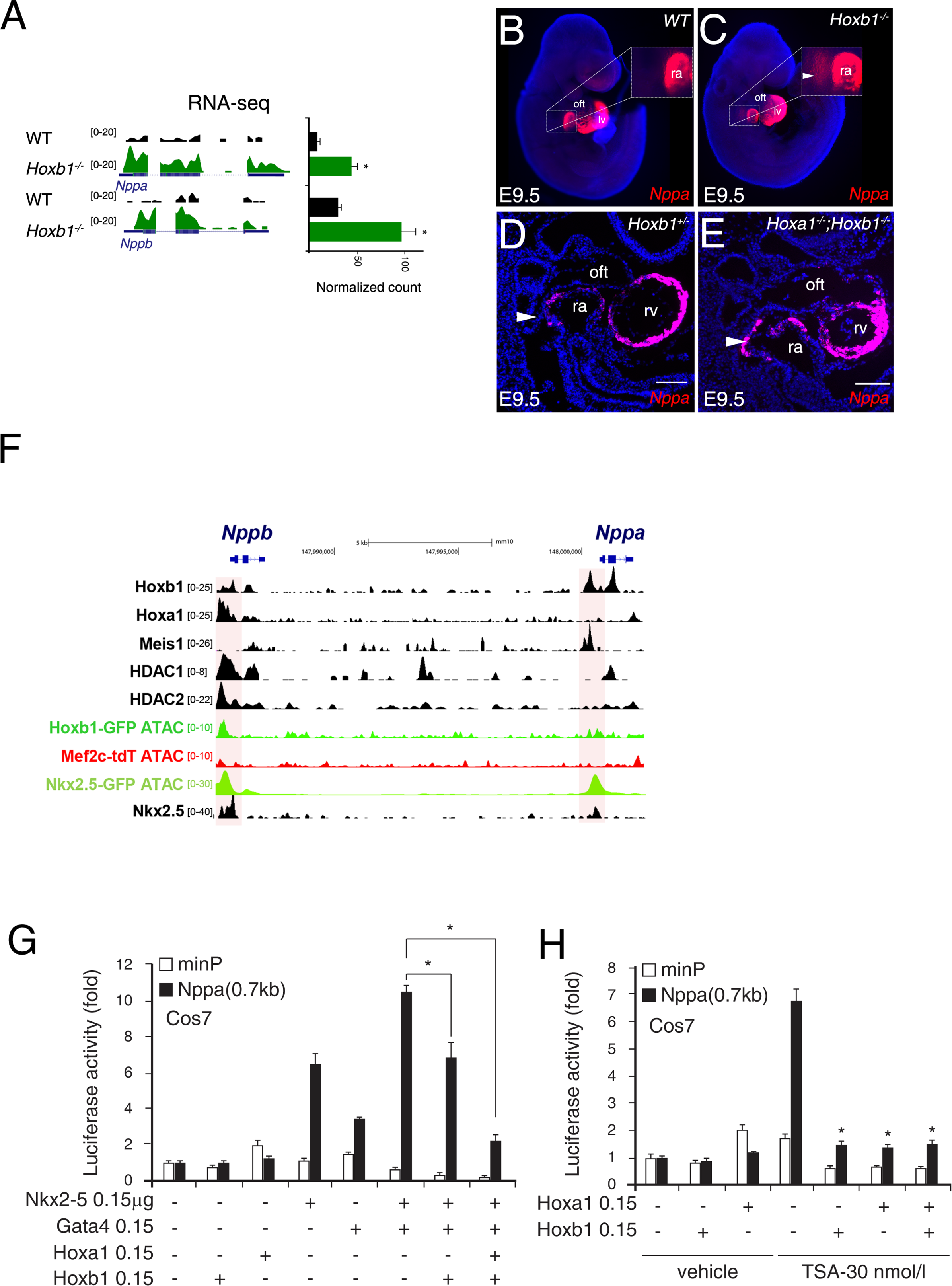
Hoxb1 regulates cardiac differentiation through transcriptional repression of myocardial genes. (A) Browser views of *Nppa* and *Nppb* gene loci with RNA-seq profiles of *Hoxb1^-/-^* (green) and wild-type (WT) population (black). Data represent the union of technical and biological replicates for each cell type. The *y*-axis scales range from 0–5 in normalized arbitrary units. (B,C) Whole mount RNA-FISH for *Nppa* in WT (B) and *Hoxb1^-/-^* (C) E9.5 embryos. Inset displays higher magnification of the posterior heart region. Nuclei are stained with Dapi. RNA-FISH on sagittal sections for *Nppa* in *Hoxb1^+/-^* (D) and *Hoxa1*^-/-^;*Hoxb1^-/-^* (E) at stage E9.5. (F) Browser view of *Nppa*, *Nppb* locus with ATAC-seq on purified cardiac cells and ChIP-seq profiles of Hoxa1 (De Kumar et al., 2017), Meis1 (Losa et al., 2017), Nkx2-5 (van den Boogaard et al., 2012), HDAC1 and HDAC2 (Whyte et al., 2012). (G) Constructs were co-transfected with Nkx2-5, Gata4, Hoxa1 and Hoxb1 expression vectors into Cos-7 cells. Luciferase activity was determined and normalized as fold over the reporter alone (mean ± SEM, n=3, *p<0.05 for Nkx2-5 and Hoxb1 *versus* Nkx2-5, using ANOVA). (H) Luciferase reporter assays on the −633/+87-bp region of the *Nppa* promoter. Cos-7 cells co-transfected with Hoxa1 and Hoxb1 expressing vector or not were treated in the absence or presence of 30 nmol/l TSA. Bars represent mean ± SEM (n=3). Statistical test was conducted using ANOVA (*p<0.05 for Hoxa1, Hoxb1 and TSA treatment versus Hoxb1 or TSA treatment). oft, outflow tract; lv, left ventricle; ra, right atrium.

## DISCUSSION

In this study, we characterize the transcriptional profile of subpopulations of SHF progenitor cells contributing to the forming heart and identify central roles of Hoxb1 in the posterior SHF. We report that forced activation of *Hoxb1* in the *Mef2c-AHF* lineage results in a hypoplastic right ventricle and show that Hoxb1 has a dual role in activating the posterior program of the SHF and inhibiting premature cardiac progenitor differentiation through the transcriptional repression of myocardial genes. Thus Hoxb1 coordinates patterning and deployment of SHF cells during heart tube elongation and altered Hoxb1 expression contributes to CHD affecting both poles of the heart tube.

### Transcriptional profile and chromatin accessibility mapping of the posterior SHF

Here, we present the first analysis of open chromatin in purified SHF progenitor cells using the ATAC-seq method. Such datasets are important to understand the tightly regulated genetic networks governing heart development and to determine how these networks become deregulated in CHDs. Integration with transcriptomes of sorted cardiac progenitor cells led us to the discovery of novel pSHF markers illustrating how this study provides a large dataset and multiple new avenues of investigation for the future. Among genes enriched in the *Hoxb1*-expressing progenitors we found several genes known to be important for the inflow tract development, including *Tbx5*, *Osr1*, *Foxf1*, *Bmp4, Wnt2*, and *Gata4*. Importantly, *Tbx5*, *Osr1*, *Foxf1*, and *Bmp4,* also contribute to the DMP, which derives from both *Hoxb1* and *Mef2c-AHF*-lineages, (Zhou et al., 2017; Burns et al., 2016; Zhou et al., 2015; Hoffmann et al., 2014; Briggs et al., 2013; Xie et al., 2012) indicating that the transcriptional expression level of these genes was highly enriched in the GFP+ relative to Tomato+ cells. Our analysis of the double positive *Hoxb1^GFP^* (GFP+) and *Mef2c-Cre;Rosa^tdT^* (Tomato+) population at E9.5 showed an activation of *Osr1* and *Adlh1a2* transcripts, whereas both *Hoxb1* and *Tbx5* transcripts are weakly expressed (Figure 2K). These results suggest a progressive change in cell identity during SHF development.

### Hox genes are required for atrioventricular septation

Although recent single-cell RNA-seq analyses identified specific pSHF and aSHF clusters, the subpulmonary and DMP specific sub-populations have not yet been characterized, probably due to the restricted number of these progenitors (Hoffmann et al., 2014;de Soysa et al., 2019; Pijuan-Sala et al., 2019). The DMP protrudes into the atrial lumen to contribute to the atrioventricular mesenchymal complex and muscular base of the atrial septum. Perturbation of DMP development in embryos mutant for transcription factors and signaling molecules regulating posterior SHF deployment results in AVSDs (De Bono et al., 2018; Rana et al., 2014; Briggs et al., 2012; Xie et al., 2012; Goddeeris et al., 2008). We observed that either *Hoxb1* or *Hoxa1;Hoxb1* loss of function leads to premature differentiation in the SHF. Importantly, premature myocardial differentiation in the SHF is known to contribute to defective DMP development and leads to AVSD (Goddeeris et al., 2008). Consistent with these observations we found AVSD with absence of the DMP in *Hoxa1^-/-^;Hoxb1^-/-^* mutant embryos, identifying Hox genes as upstream players in the etiology of this common form of CHD. Among the pSHF genes upregulated in *Hoxb1^GoF^;Mef2c-Cre* and downregulated in *Hoxb1^-/-^* mutant embryos were *Bmp4* and *Gata4*, suggesting that Hoxb1 may directly activate these genes (Figure 5). Bmp4 is expressed in the DMP (Briggs et al., 2016; Burns et al., 2016; Sun et al., 2015; Briggs et al., 2013) and mutations in BMP4 and the BMP receptor Alk2 have been implicated in atrial septal defects and AVSDs (Smith et al., 2011). Gata4 is required for atrioventricular septation (Liao et al 2008, Zhou et al 2017).

### Hoxb1 is required to repress cardiac differentiation in the pSHF

Examination of ectopic expression of *Hoxb1* in the *Mef2c-Cre* derivatives showed a hypoplastic right ventricle phenotype. RNA-seq analysis showed that ectopic expression of *Hoxb1* results in mis-specification of the anterior program leading to the downregulation of genes involved in cardiac differentiation. Consistent results were observed in *Hoxb1^-/-^* mutant embryos where GO analysis showed upregulated genes related to “cardiac muscle tissue development” and “muscle system process”. This aberrant gene program prompted us to hypothesize that Hoxb1 blocks the differentiation process by inhibiting activation of a set of myocardial genes. Our observations in the mES cell system confirmed the repressive function of Hoxb1 during cardiac differentiation. *In vitro* analysis in conjunction with our RNA-seq analysis confirmed that Hoxb1 directly represses the activation of structural myocardial genes. Hoxb1 activity on *Nppa* promoter was consistent with our *in vivo* analysis. The 0.7-kb *Nppa* fragment is responsible for the developmental expression pattern of the *Nppa* gene and is itself regulated by Gata4 and Nkx2-5 (Habets et al., 2002). We provide molecular evidence that Hoxb1 functions to repress differentiation in the pSHF. Several studies have suggested that Hoxb1 plays a repressive role in differentiation of other cell types (Chen et al., 2012; Bami et al., 2011). Indeed, HOX transcription factors can function as activators or repressors (Mann et al., 2009; Saleh et al., 2000) raising the question of how gene activation versus repression is determined. Our analysis of ATAC-seq in pSHF population demonstrated overrepresentation of specific transcription factor motifs. Several of these transcription factors, including Nkx2-5, Gata4, Pbx1/2/3 and Meis1/2 have been previously associated with cardiac differentiation. Nkx2-5 cooperates with Hox factors to regulate the timing of cardiac mesoderm differentiation (Behrens et al., 2013). TEAD proteins regulate a variety of processes including cell proliferation survival and heart growth (Lin et al., 2016; von Gise et al., 2012). Unexpectedly, our results implicate TEAD proteins in activating regulatory genes and potentially promoting the proliferation/outgrowth of cardiac progenitors in the pSHF (Francou et al., 2017). The Hox protein family has been previously shown to interact other with homeodomain proteins, including the TALE-class Pbx and Meis family members (Lescroart and Zaffran, 2018). This further supports the conclusion that Hoxb1 uses cofactors to activate or repress cardiac genes. Repression is mediated by direct recruitment of repressor complexes, such as NuRD, or through maintenance of repressed chromatin states, such as those mediated by Polycomb complexes functioning in an HDAC complex (Schuettengruber and Cavalli, 2009). Several Hox proteins reportedly bind HAT or HDACs enzymes (Ladam and Sagerstrom, 2014; Saleh et al., 2000). Meis proteins promote HATs recruitment by displacing HDACs to permit HAT binding (Shen et al., 2001). Our results suggest that Meis proteins might thus modulate HAT/HDAC accessibility at Hox-regulated regulatory sequences to delay differentiation in the SHF. Indeed, a recent study using live imaging of cell lineage tracing and differentiation status suggests that in mouse a discrete temporal lag can be observed between the first and second waves of differentiation that form the mouse heart (Ivanovitch et al., 2017). Such a delay of differentiation is essential to orchestrate early cardiac morphogenesis. Hoxb1 may thus contribute to controlling this differentiation delay, in particular through maintaining pSHF progenitor cells in an undifferentiated state until they are added to the venous pole or the inferior wall of the outflow tract. Future work will define how *Hoxb1* expression is downregulated to release the cardiac differentiation process during SHF deployment.

## METHODS

### Mice

All animal procedures were carried out under protocols approved by a national appointed ethical committee for animal experimentation (Ministère de l’Education Nationale, de l’Enseignement Supérieur et de la Recherche; Authorization N°32-08102012). The *Hoxb1^GFP^, Hoxb1^IRES-Cre^* alleles and the *Tg*(*CAG-Hoxb1-EGFP)^1Sza^* transgene (*Hoxb1^-^* and *Hoxb1^GoF^* respectively) have been previously described (Zaffran et al., 2018; Gaufo et al., 2000). The reporter lines *Gt(ROSA)26Sor^tm1Sor^* (*R26R*), *Gt(ROSA)26Sor^tm9(CAG-tdTomato)Hze^ (Rosa^tdTomato^)* have been previously described (Madisen et al., 2010; Soriano, 1999). The *Mef2c-AHF-Cre (Mef2c-Cre), Mlc1v-nlacZ-24* and *Mlc3f-nlacZ-2E* mice have been previously described (Verzi et al., 2005; Zaffran et al., 2004; Kelly et al., 2001). *Hoxb1^GoF/+^* mice were maintained on a C57Bl/6 background and inter-crossed with *Mef2c-Cre* with or without *Mlc1v24* to generate compound heterozygous embryos at a Mendelian ratio.

### Cell Culture

Mouse ES^Tet-On/Hoxb1^ ES lines were generated by the Gavalas laboratory (Gouti and Gavalas, 2008). ES cells were cultured on primary mouse embryonic fibroblast feeder cells. ES cells medium was prepared by supplementing GMEM-BHK-21 (Gibco) with 7.5% FBS, 1% non-essential amino acids, 0.1 mM beta-mercaptoethanoland LIF conditioned medium obtained from pre-confluent 740 LIF-D cells that are stably transfected with a plasmid encoding LIF (Zeineddine et al., 2006). For cardiac differentiation, ES cells were re-suspended at 25=10^4^ cells/ml in GMEM medium supplemented with 20 % fetal calf serum, 1% non-essential amino-acids, and 0.1 mM beta-mercaptoethanol in hanging drops (22 μl) plated on the inside lids of bacteriological dishes. After 48hrs EBs were transferred in 10 ml medium to 10 cm bacteriological dishes. At day 5 EBs were plated on tissue culture dishes coated with gelatin, allowed to adhere. Expression of Hoxb1 was induced by addition of doxycycline (DOX) (Sigma −1 or 0.2μg/ml) from day 4 to the end of the experiment. The medium was changed every two days.

### Cell Sorting

E9.5 (16s) transgenic progenitor heart regions were dissected, pooled (n>3 embryos for each genotype) and digested with 0.25% Trypsin/EDTA (Invitrogen), neutralized in DMEM (Invitrogen) containing 5% FBS and 10 mmol/L HEPES (Invitrogen), rinsed and resuspended in PBS, and passed through a 70-mm nylon cell strainer (Falcon). Samples were sorted on a FacsAria flow cytometer (BD) using FACSDiva 8.0.1 software. Samples were gated to exclude debris and cell clumps. The number of E9.5 *Hoxb1^GFP^* progenitor cells and *Mef2c-Cre;Rosa^tdT^* progenitor cells per embryo obtained were typically 600 to 900, respectively. Fluorescent cells were collected into PBS and processed for RNA extraction or ATAC-seq.

### Histological and Immunostaining

Standard histological procedures were used (Roux et al., 2015). Heart from *Hoxb1^GoF^;Mef2c-Cre* and littermate controls were fixed in neutral-buffered 4% paraformaldehyde in PBS, rinsed, dehydrated, paraffin-embedded and tissue sections cut at 8μm. Sections were stained with Harris’ hematoxylin and eosin (H&E) (Sigma). For immunostaining embryos from *Hoxb1^-/-^* or *Hoxb1^GoF^;Mef2c-Cre* and littermate controls were fixed at 4°C for 20min in 4% paraformaldehyde, rinsed in PBS, equilibrated to 15% sucrose and embedded in O.C.T. Cryo-sections were cut at 12μm, washed in PBS and pre-incubated in blocking solution (1%BSA, 1% Serum, 0.2% Tween20 in PBS). Primary antibodies were applied overnight at 4°C, followed by secondary detection using Alexa Fluor conjugated (Molecular Probes) secondary antibodies. Sections were photographed using an AxioImager Z2 microscope (Zeiss) and photographed with an Axiocam digital camera (Zen 2011, Zeiss). The following primary antibodies were used in this study: rabbit anti-Hoxb1 (Covance; 1/200), rabbit anti-GFP (Invitrogen; 1/500), mouse anti-αactinin (sigma; 1/500), mouse anti-MF-20 (DHSB; 1/100), Rabbit anti-Caspase3 (Cell Signaling Technology, 1/300), rabbit anti-phospho-Histone H3 (Merck; 1/400), and mouse anti-Islet1 (DSHB; 1/100), rabbit anti-Tbx1 (Lifescience Ls-C31179, 1/100), goat anti-Tbx5 (Santa Cruz sc-7866, 1/250).

### X-gal staining

X-gal staining was performed on whole-mount embryos as described previously (Roux et al., 2015). For each experiment, a minimum of three embryos of each genotype was observed. Embryos were examined using an AxioZoom.V16 microscope (Zeiss) and photographed with an Axiocam digital camera (Zen 2011, Zeiss).

### *In Situ* hybridization

RNA-FISH was performed according to the protocol of the RNAscope Multiplex Fluorescent v2 Assay (cat. no.323110), which detects single mRNA molecules. In briefly, E8.5 and E9.5 embryos were fixed for 20-30h in 4% paraformaldehyde and then dehydrated in methanol. Whole-mount RNA-FISH was performed as previously described (de Soysa et al., 2019). Embryos were imaged using an AxioZoom.V16 microscope (Zeiss). The following probes were used: mm-*Hoxb1*-C1 (cat no. 541861), mm-*Bmp4*-C1 (cat no. 401301), *mm-Gata4* (cat no.), *mmAldh1a2* (cat no.) mm-e*GFP*-C3 (cat no. 400281), mm-*tdTomato*-C3 (cat no. 317041), mm-Tbx1-C1 (cat no. 481911), mm-*Tbx5*-C2 (cat no. 519581), and mm-*Osr1*-C2 (cat no. 496281-C2).

### ATAC-seq

For each sample, 10,000 FACS-sorted cells were used (n>3 embryos for each genotype). Cell preparation, transposition reaction, and library amplification were performed as previously described (Buenrostro et al., 2013). Paired-end deep sequencing was performed using a service from GenomEast platform (IGBMC, Strasbourg).

### Processing of ATAC-sequencing data and statistical analysis

Raw ATAC-Seq reads were aligned with the SNAP aligner (http://snap.cs.berkeley.edu/) on the reference GRCm38 mouse genome. Deduplicated reads were marked and, following ENCODE specifications (https://www.encodeproject.org/atac-seq/), unmapped, not primarily aligned, failing platform and duplicated reads were removed using samtools (-F 1804). Properly paired reads were kept (samtools -f 2). Finally, reads mapping blacklisted regions for mouse genome mm10 provided by ENCODE (Carroll et al., 2014) were excluded from the analysis.

To evaluate reproducibility between replicates and retain peaks with high rank consistency, we applied the Irreproducible Discovery Rate (IDR; https://f1000.com/work) methodology from ENCODE. Only peaks with an IDR value lower than 0.1 were retained.

Narrow peaks were called with MACS 2.1.1. (https://f1000.com/work) BigWig files were generated from bedGraph files to visualize fold enrichment and p-value for all regions within UCSC genome browser.

Differential

A MA plot (log_2_ fold change vs. mean average) was used to visualize changes in chromatin accessibility for all peaks. MA plot depicts the differences between ATAC-seq peaks in the experimental samples by transforming the data onto M (log ratio) and A (mean average) scales, then plotting these values. Differential chromatin accessibility is expressed as a log fold change of at least 2 folds and a P-value of <0.1 and reveals the relative gain of chromatin regions in GFP+ cells (above the 0 threshold line) as compared to the gain in the Tomato+ cells (below the 0 threshold line).

Differential peaks between GFP+ (*Hoxb1-GFP*) and Tomato+ (*Mef2c-Cre;Rosa^tdT^*) samples were identified using a bed file containing selected peaks from IDR methodology and the DiffBind R package (10.18129/B9.bioc.DiffBind). Peaks with FDR (False Discovery Rate) at 10% were kept.

Differential peaks were annotated using the Homer^7^ software and genes in the vicinity of peaks (+/- 150-kb from summit) were selected.

In order to perform motif analysis, we generated a Fasta file containing all sequences surrounding peak summits (+/- 200-bp) and used the Homer findMotifsGenome feature. Known and *de novo* motifs were identified. All possible order-3 combinations motifs were generated for each known and *de novo* motifs. To assess enrichment of motifs combinations, *p*-values were computed using 20,000 random sequences of 400-bp from the mouse genome. Fisher’s exact test was applied to compare random and peaks sequences.

### RNA-seq

Total RNA was isolated from the pharyngeal region and sorted cells with NucleoSpin RNA XS (Bioke) following the protocol of the manufacturer. cDNA was generated and amplified with the Ovation RNAseq v2 kit (NuGEN). Briefly, 2 ng of total RNA were used for mixed random-/polyA-primed first-strand cDNA synthesis. After second strand synthesis, the double-stranded cDNA was amplified by single primer isothermal amplification, and the amplified cDNA was bead-purified (AmpureXP, Beckman-Coulter). Paired-end deep-transcriptome sequencing was performed using a service from GenomEast platform (IGBMC, Strasbourg).

E9.5 *Mef2c-Cre:Rosa^tdTomato^* and *Hoxb1^GFP^* embryos were dissected on ice-cold PBS to isolate the SHF and cells were FACS. After FACS Tomato+ and Tomato-as well as GFP+ and GFP-cells were homogenized in Trizol (Invitrogen) using a Tissue-lyzer (Qiagen). RNA was isolated from pharyngeal region of E8.5 wild-type and *Hoxb1^-/-^*, and E9.5 control and *Hoxb1^GoF^;Mef2c-Cre* embryos (n=3 embryos per each genotype). RNAs of each genotype were pooled to obtain one replicate. RNA was prepared using the standard Illumina TrueSeq RNASeq library preparation kit. Libraries were sequenced in a Hiseq Illumina sequencer using a 50-bp single end elongation protocol. For details of analyses of RNA-seq data see Supplemental file 1. Resulting reads were aligned and gene expression quantified using RSEM v1.2.3 (Li and Dewey, 2011) over mouse reference GRCm38 and Ensembl genebuild 70. Gene differential expression was analyzed using EdgeR R package (McCarthy et al., 2012; Robinson et al., 2010). Genes showing altered expression with adjusted P<0.05 were considered differentially expressed. For the set of differentially expressed genes a functional analysis was performed using Ingenuity Pathway Analysis Software and DAVID (Huang da et al., 2009), and some of the enriched processes were selected according to relevant criteria related to the biological process studied.

### RNA-Seq data processing

Raw RNA-seq reads were aligned using the STAR aligner version 2.5.2 (Dobin et al., 2013) on the reference GRCm38 mouse genome. Coverage visualization files (WIG) were generated with the STAR aligner software and were converted into BigWig files using UCSC wigToBigWig files to allow their visualization within the UCSC genome Browser.

In parallel, transcripts abundance was computed using HTSEQ-count 0.9.1 (Anders et al., 2015) and the background was estimated through read counts from intergenic regions using windows of 5-kb length.

Normalization and differential gene expression analysis between conditions were performed using R (R version 3.3.4) and DESEQ2 (Love et al., 2014).

For each sample, genes with null expression were removed and we set the 95th percentile of the intergenic read counts as the threshold of detection (log_2_(normalized count + 1)). Heatmap were generated with the Pheatmap R package.

### Functional Annotation

For RNA-seq and ATAC-seq, genes lists were annotated with the ClusterProfiler (Yu et al., 2012) system. Circos plots were generated with Goplot package (Walter et al., 2015).

### *In vitro* reporter assays

Luciferase reporter constructs were co-transfected with expression constructs for human *HOXB1*, *GATA4* (Stefanovic et al., 2014; Singh et al., 2009) and *NKX2-5* (Singh et al., 2009). Constructs were transfected into Cos-7 cells with the PEI transfection reagent. Cell extracts and luciferase assays were performed following the protocol of the manufacturer (Promega). Mean luciferase activities and standard deviations were plotted as fold activation when compared with the empty expression plasmid.

### Quantitative RT-PCR analysis

Total RNA was isolated from pharyngeal regions and sorted cells with NucleoSpin RNA XS (Bioke) following the protocol of the manufacturer. cDNA was generated using the AffinityScript Multiple Temperature cDNA synthesis kit (Agilent). The expression level of different genes was assessed with quantitative real-time PCR using the LightCycler Real-Time PCR system (Roche Diagnostics) and the primers described in the Supplementary data. Values were normalized to HPRT expression levels.

### Statistics

Statistical analyses were performed using unpaired two-tailed t-test to assess differences between two groups. Data are presented as mean ± SD. A P value of <0.05 was considered significant.

## ADDITIONAL INFORMATIONS

### Accession codes

Sequencing data have been deposited in Gene Expression Omnibus under accession number GSE123765 (ATAC-seq on GFP+ and Tomato+ cells); GSE123771 (RNA-seq on GFP+ and Tomato+ cells); GSE123772 (RNA-seq on *Hoxb1^Go F^* vs. control embryos) and GSE123773 (RNA-seq on *Hoxb 1^-/-^* vs. wild-type embryos).

## ACKNOWLEDGEMENTS

We are grateful to Heather Etchevers for her comments on the manuscript. We thank the animal facility, particularly Adeline Gatha, and Françoise Mallet for the cell sorting (INSERM U1068 / CNRS UMR 7258). Sequencing was performed by the IGBMC Microarray and Sequencing platform. This work was supported by the FG National Infrastructure, funded as part of the “Investissements d’Avenir” program managed by the Agence Nationale pour la Recherche (ANR-10-INBS-0009). We thank the Leducq Foundation for funding the program “Imaging cardiac cell lineages at the origin of congenital heart disease” in the frame of the Research Equipment and Technological Platform Awards (Coordinator M.P. and Partners R.K, S.Z.). This work was supported by the INSERM, the Agence National pour la Recherche (Transcardiac and Heartbox projects, S.Z. and R.K.), the Association Française contre les Myopathies (S.Z.), and the Fondation pour la Recherche Medicale DEQ20150331717 (R.K.). S.S. was supported by post-doctoral awards from l’Institut de France Lefoulon-Delalande and H2020-MSCA-IF-2014. B.L. was supported by a post-doctoral fellowship from the Fondation pour la Recherche Médicale.

## AUTHOR CONTRIBUTIONS

S.S., B.L., JP. D, F.L., L.A., C.M-Z., D.S., performed and analyzed the experiments. C.B., A.G., M.P., R.K. and S.Z. reviewed the data. S.S., B.L., F.L., and S.Z. designed experiments, and wrote the first draft of the manuscript, which was finalized upon discussion with all authors.

## Supplemental Figure Legends

**Figure 2 – Figure supplement 1:**
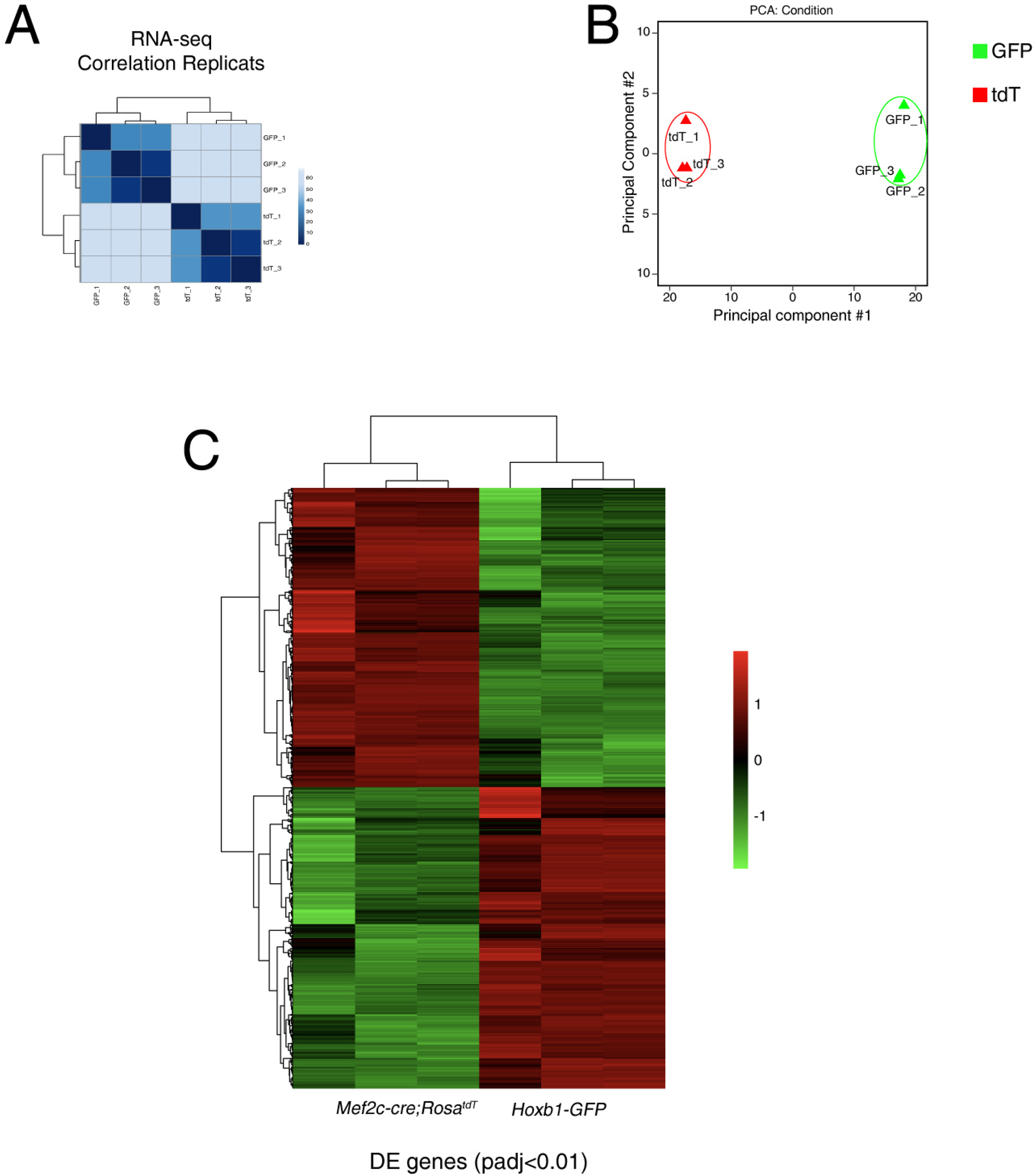
Quality assessment of RNA-seq data performed with purified cardiac progenitor cells. (A) Replicate correlation for RNA-seq datasets from GFP+ and Tomato+ progenitor cells. (B) Principal component analysis (PCA) of RNA-sequencing datasets from GFP+ and Tomato+ progenitor cells. (C) Unsupervised hierarchical clustering of all differentially expressed genes between GFP+ and Tomato+ progenitor cells.

**Figure 2 – Figure supplement 2:**
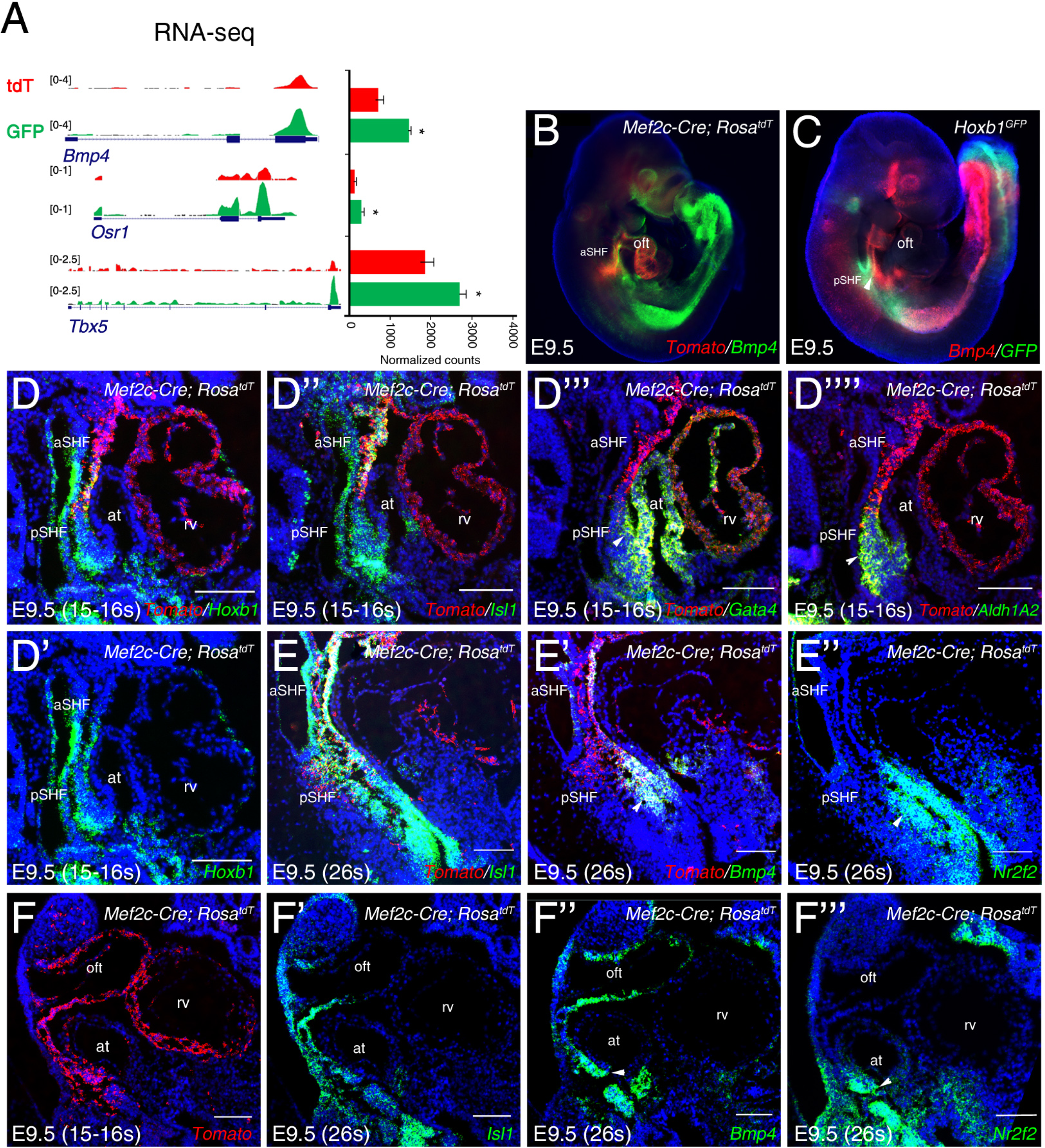
Spatial validation of marker gene expression in cardiac progenitor populations. (A) RNA-sequencing datasets visualized for the pSHF markers *Tbx5*, *Osr1* and *Bmp4.* (B) Whole-mount RNA-FISH analysis showing the distribution of *Bmp4* transcript (green) in cells overlapping with the posterior border of *Mef2c-Cre* lineage contribution (red). (C) *Bmp4* (red) is expressed in the pSHF overlapping with *Hoxb1-GFP* (green). (D-F) RNA-FISH analysis on serial sagittal sections of *Mef2c-Cre;Rosa^tdT^* embryos, showing expression of genes enriched in the GFP+ progenitor population (E and F are from the same embryo). *Hoxb1*, *Gata4*, *Aldh1a2* and *Nr2f2* mark the pSHF, whereas *Bmp4* expression is enriched in the pSHF compared to the aSHF. Scale bars: 100 μm.

**Figure 3 – Figure supplement 3:**
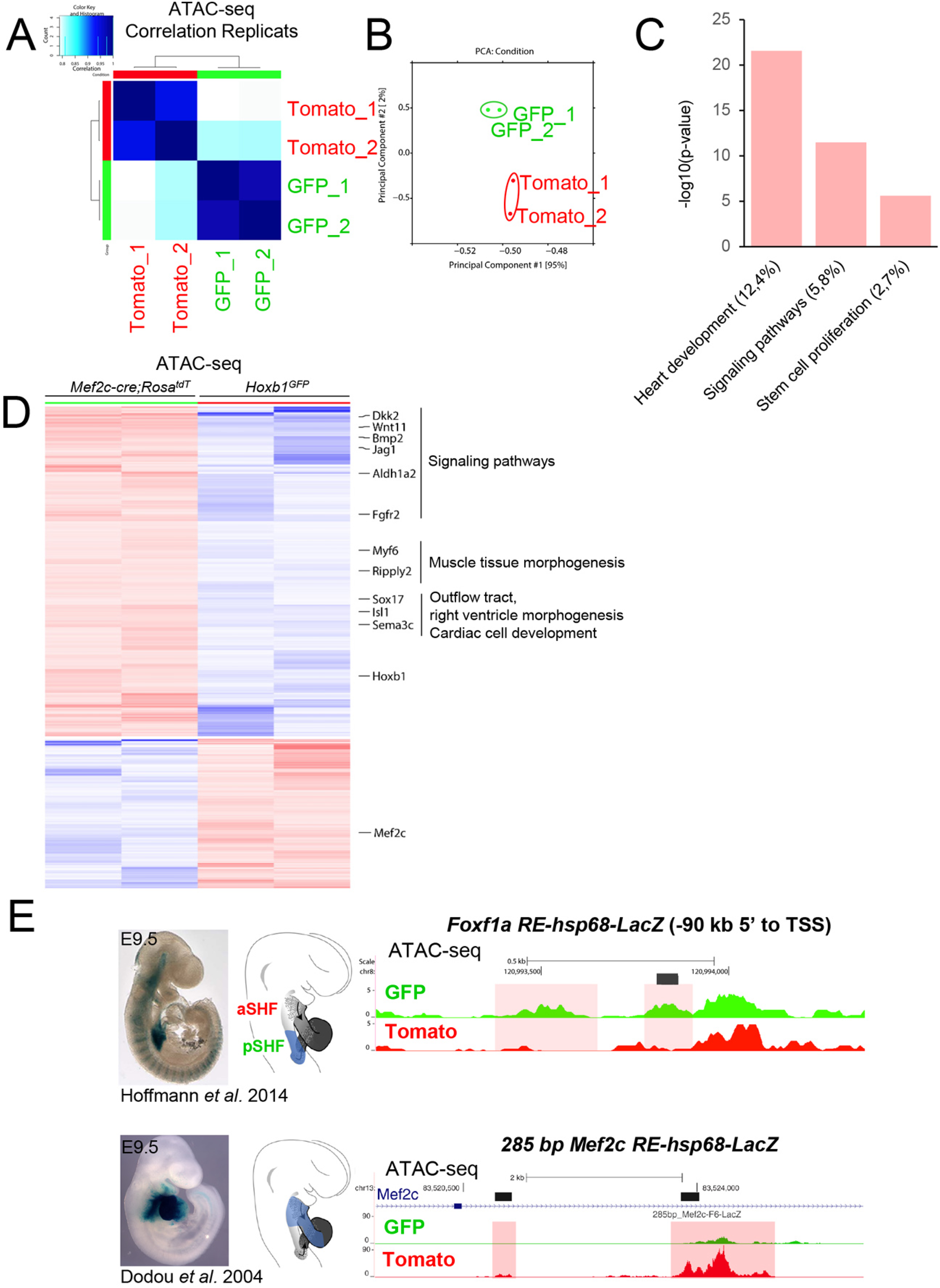
Quality assessment of ATAC-seq data performed with purified cardiac progenitor cells. (A) Replicate correlation for ATAC-seq datasets from GFP+ and Tomato+ progenitor cells. (B) Principal component analysis (PCA) of ATAC-sequencing datasets from GFP+ and Tomato+ progenitor cells. (C) The Gene Ontology (GO) analysis results for the gene loci harboring peaks strictly present in the GFP+ population. (D) Heat maps show the ATAC-seq enrichment (±150 kb region upstream from the annotated TSS). (E) ATAC-sequencing on SHF cells identifies previously established SHF REs for *Foxf1a* (top) and *Mef2c* (bottom).

**Figure 4 – Figure supplement 4:**
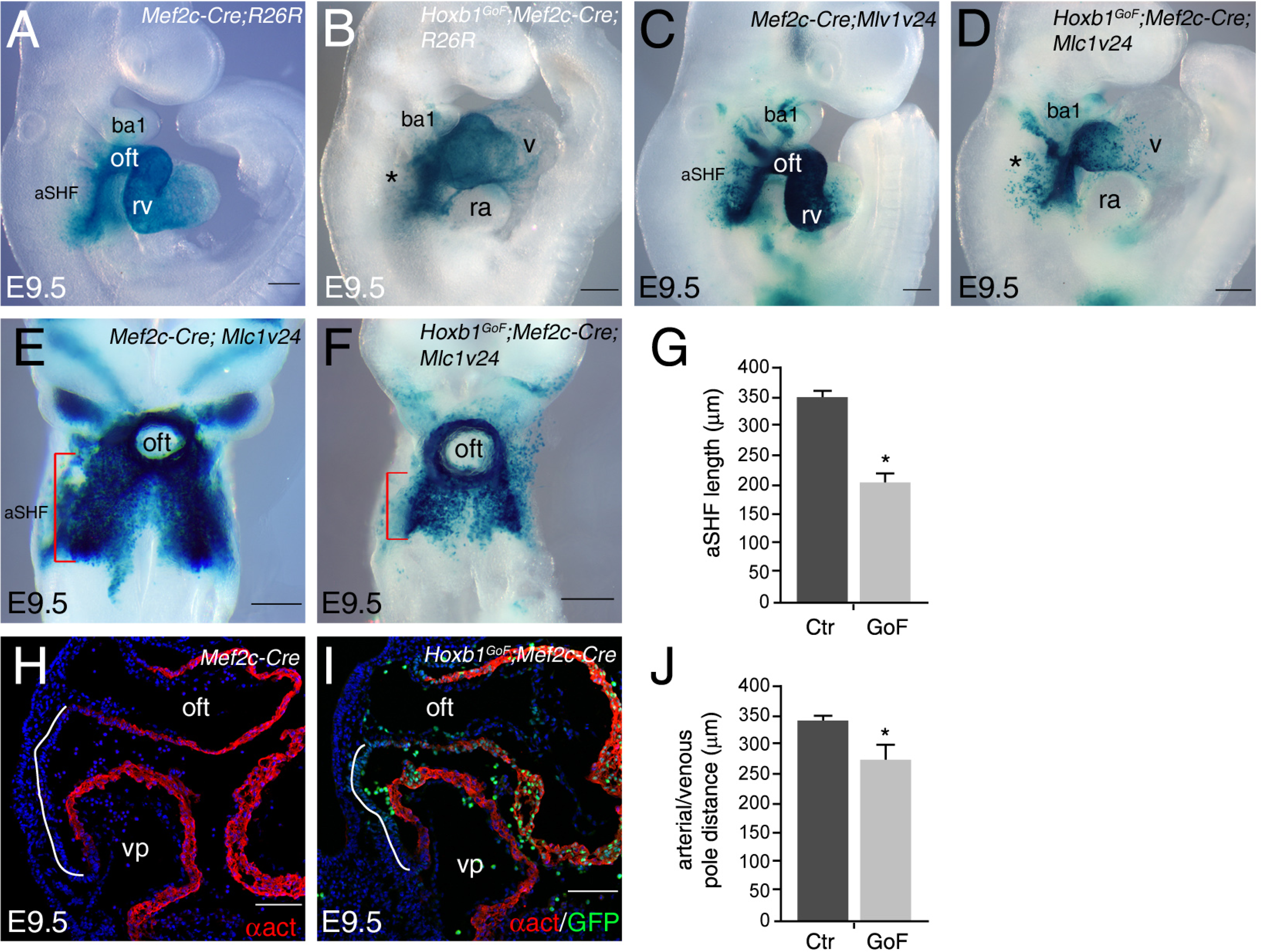
Reduction of SHF length in *Hoxb1^GoF^;Mef2c-Cre* embryos. (A-F) X-gal staining of control and *Hoxb1^GoF^* embryos carrying out *R26R* (A,B) or *Mlc1v24* (C-F) transgenes at E9.5. (A,B) Anterior SHF is disrupted in *Hoxb1^GoF^*;*Mef2c-Cre* embryos. (E,F) X-gal staining showing anterior SHF in *Mef2c-Cre*;*Mlc1v24* (control; E) and *Hoxb1^GoF^*;*Mef2c-Cre;Mlc1v24* (F) embryos at E9.5. (G) Measurement of staining revealed a decrease of anterior SHF length in *Hoxb1^GoF^*;*Mef2c-Cre;Mlc1v24* compared to control embryos. (H,I) Immunofluorescence with α-actinin (α act, red) and GFP (green) on *Mef2c-Cre* (control; H) and *Hoxb1^GoF^*;*Mef2c-Cre* (I) embryos at E9.5. (J) Measurement of distance between arterial and venous poles reveals a reduction of dorsal pericardial wall length in *Hoxb1^GoF^*;*Mef2c-Cre* embryos. All measures were calculated from n=6 embryos for each genotype. Histograms are expressed as mean ±SEM. *p* values were determined by Student’s *t* test. (\**p*<0.01). Scale bars: 100μm (H,I) and 200μm (A-D,E,F).

**Figure 5 – Figure supplement 5:**
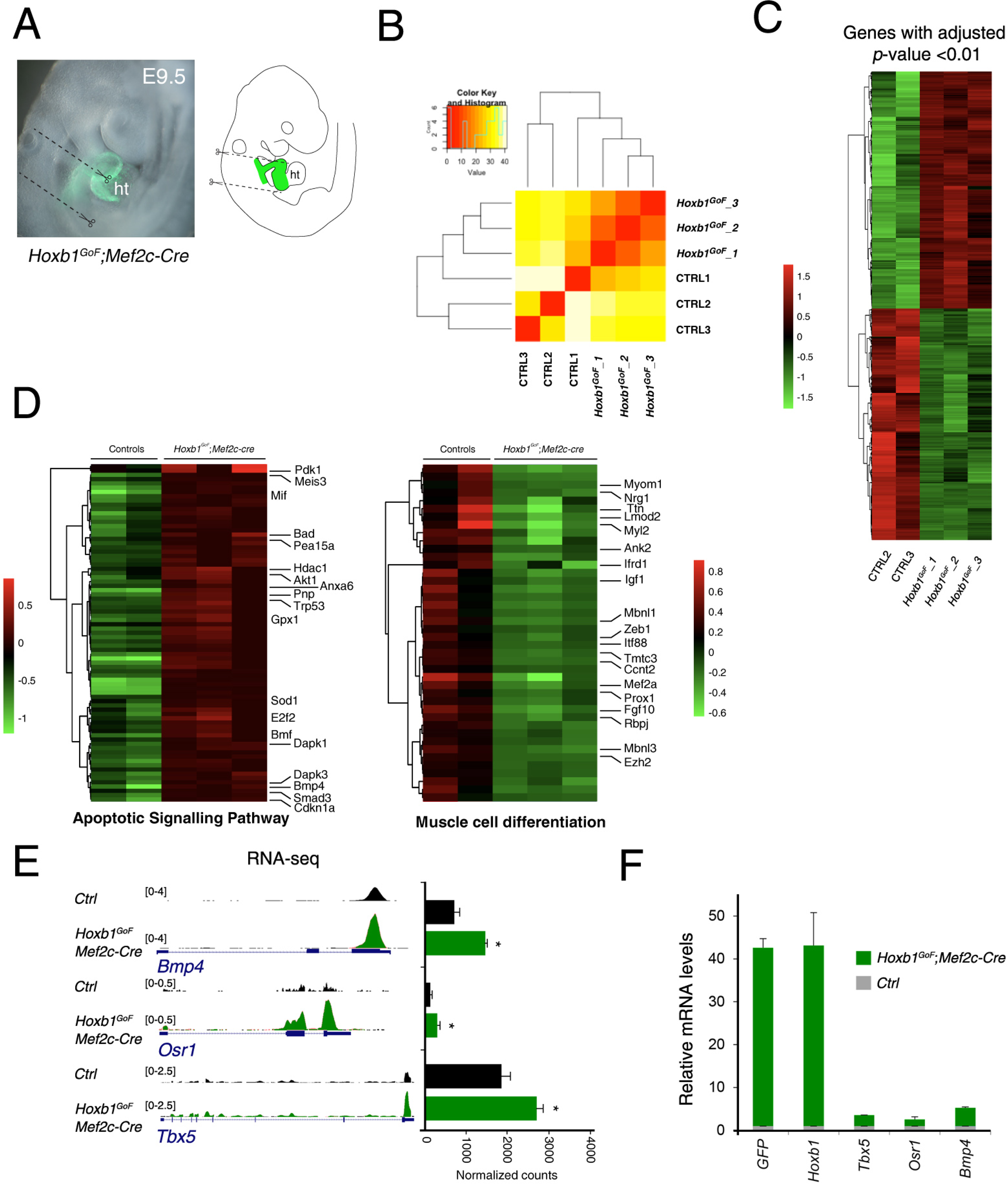
Quality assessment of RNA-seq data performed on the *Hoxb1^GoF^*;*Mef2c-Cre* embryos. (A) Macroscopic view of *Hoxb1^GoF^*;*Mef2c-Cre* embryos at E9.5 and the micro-dissected region comprising the SHF progenitors and excluding the forming heart. (B) Replicate correlation for RNA-seq datasets from *Hoxb1^GoF^*;*Mef2c-Cre* (GoF) and control (Ctrl) regions. (C) Unsupervised hierarchical clustering of all differentially expressed genes between GoF and control regions. (D) Heatmap of “apoptotic signaling pathway” and “muscle cell differentiation” associated genes analyzed by RNA-seq. (e) RNA-seq datasets visualized for the pSHF markers *Tbx5*, *Osr1* and *Bmp4*. (F) Real time PCR validation of RNA-seq results. Data were normalized to HPRT and expressed as fold increase over control samples.

**Figure 5 – Figure supplement 6:**
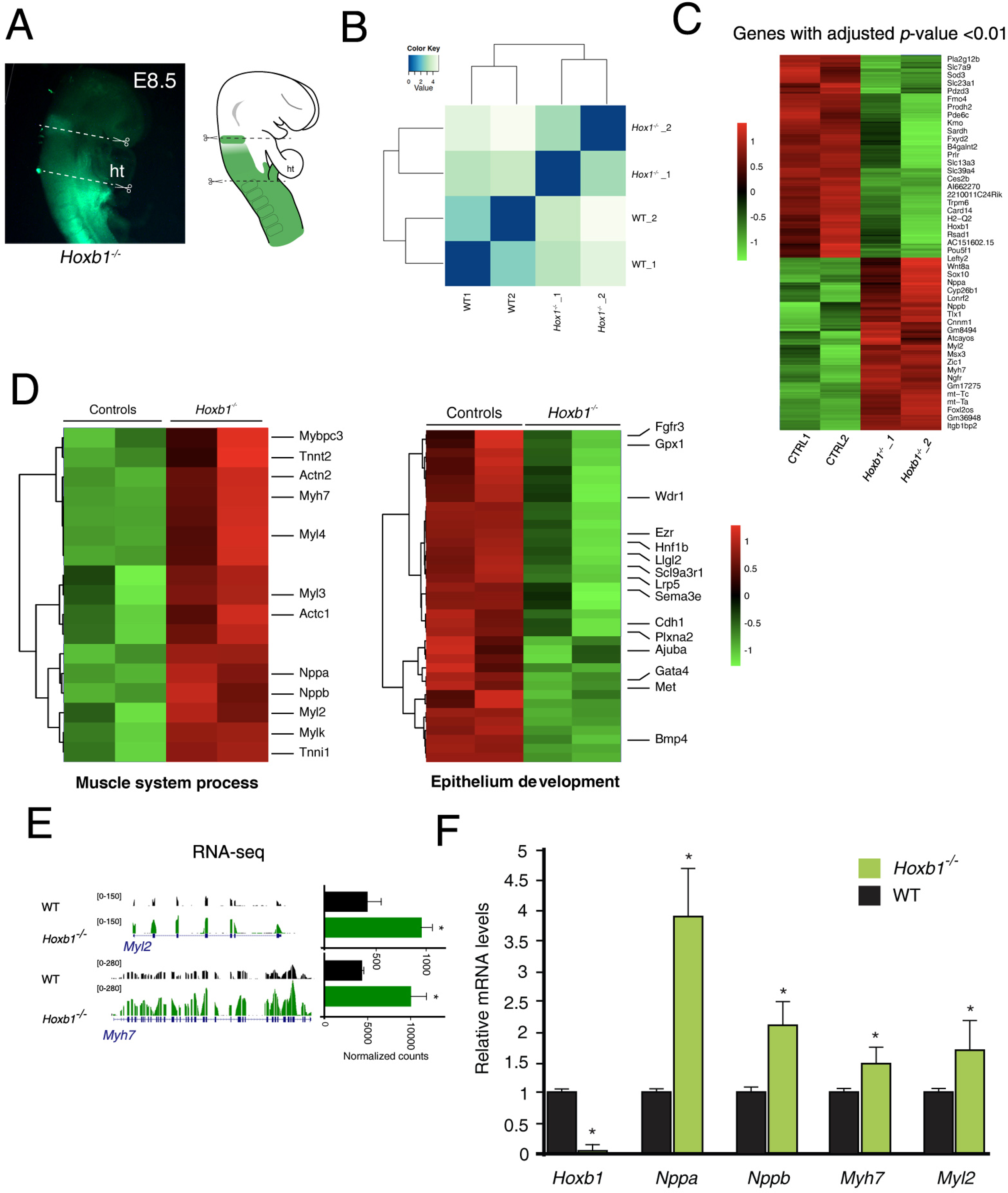
Quality assessment of RNA-seq data performed on the *Hoxb1^-/-^* embryos. (A) Macroscopic view of *Hoxb1^-/-^* embryos at E8.5 (6-8ps) and the micro-dissected region comprising the SHF progenitors. (B) Replicate correlation for RNA-seq datasets from *Hoxb1^-/-^* and wild-type (WT) regions. (C) Unsupervised hierarchical clustering of all differentially expressed genes between *Hoxb1^-/-^* over WT regions. (D) Heatmap of “muscle system process” and “epithelium development” associated genes analyzed by RNA-seq. (E) RNA-sequencing datasets visualized for the cardiac differentiation markers *Myl2* and *Myh7*. (F) Real time PCR validation of RNA-seq results. Data were normalized to HPRT and expressed as fold increase over WT samples.

**Figure 7 – Figure supplement 7:**
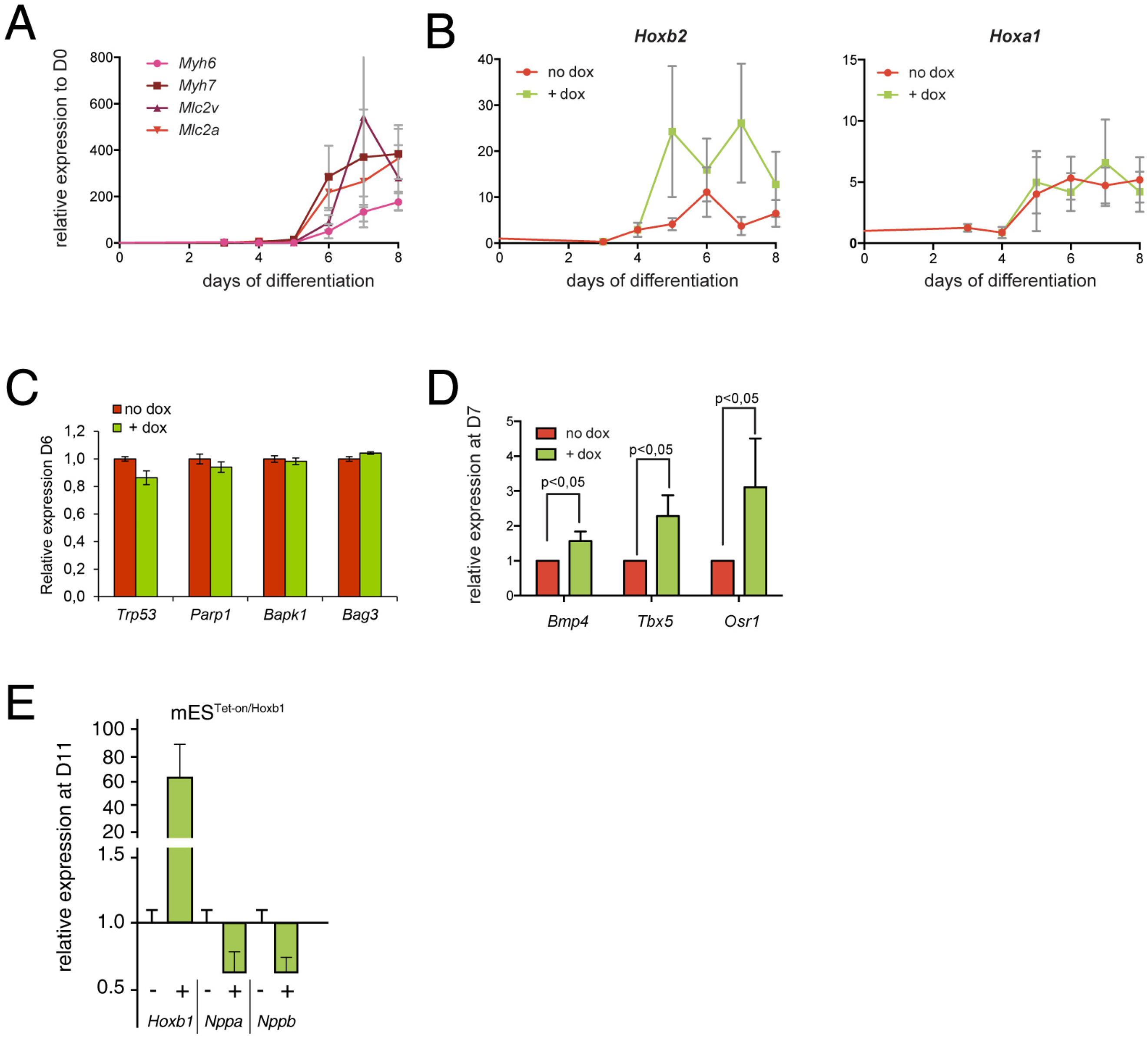
Expression analysis using the mES cells overexpressing model. (A) Kinetics of myocardial genes (*Myh6*, *Myh7*, *Myl2* (*Mlc2v*), *Mlc2a*) during mES cell differentiation as measured by real time RT-PCR. (B) Kinetics of *Hoxa1* and *Hoxb2* during mES cell differentiation after induction of *Hoxb1* expression (+ dox) or in control condition (no dox). Results are normalized for gene expression in undifferentiated mES cells (D0). (C) Relative expression of apoptosis genes (*Trp53*, *Parp1*, *Bapk1, Bag3*) at D6 of mES cell differentiation as measured by real time RT-PCR in mES cells induced or not for Hoxb1 expression (+ dox vs no dox). (D) Relative expression of posterior markers of the SHF (*Bmp4*, *Tbx5*, *Osr1*) at D7 of mES cell differentiation as measured by q-RT PCR in mES cells induced or not for Hoxb1 expression (+ dox vs no dox). (E) Relative expression of *Hoxb1*, *Nppa* and *Nppb* at D11 of mES cell differentiation as measured by real time RT-PCR in mES cells induced or not for Hoxb1 expression (+ dox vs no dox). Results are normalized for mRNA expression in untreated cells (no dox). Error bars indicate mean +/-SEM; n=4 experiments.

**Figure 7 – Figure supplement 8:**
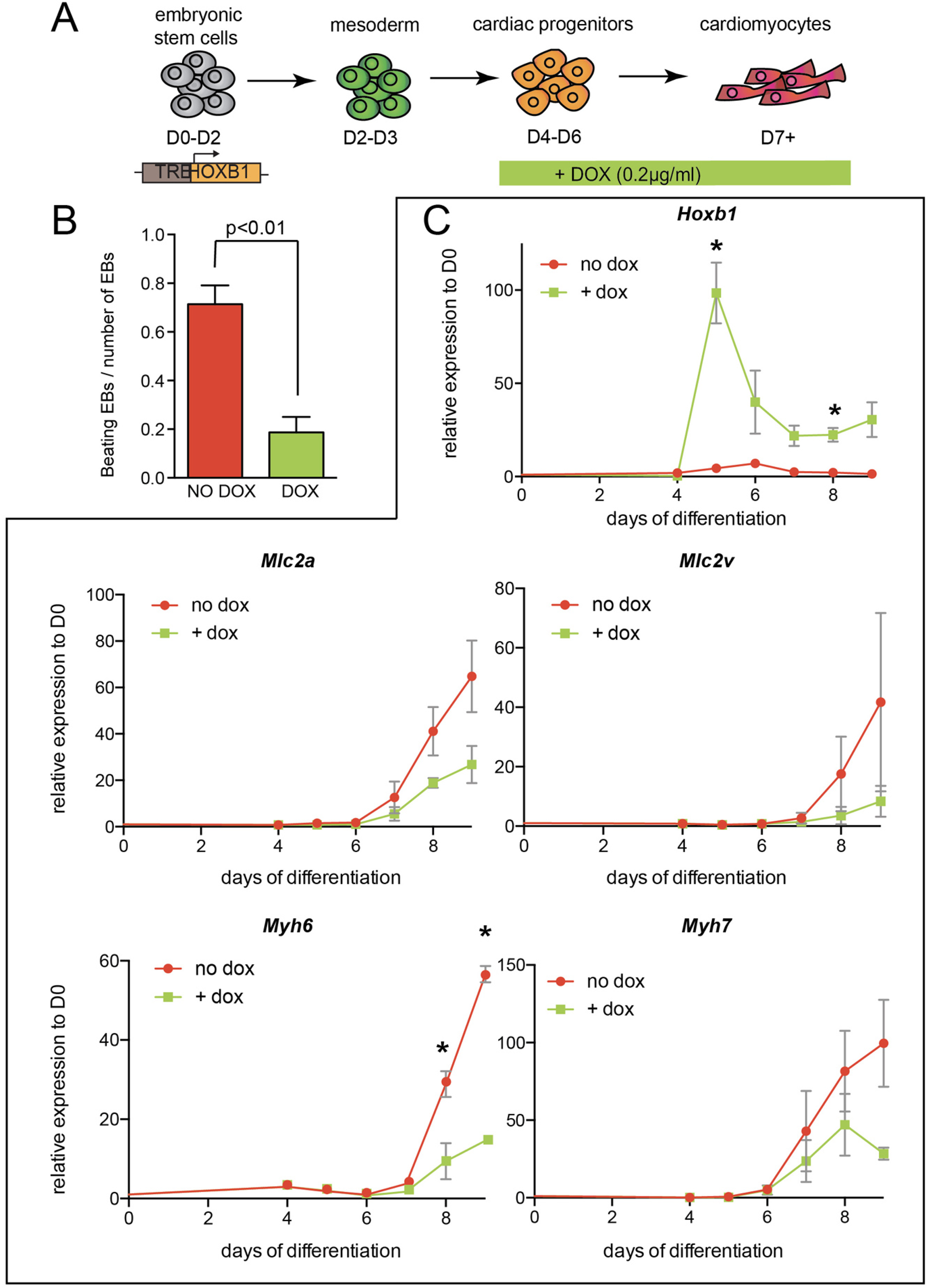
Arrested cardiac differentiation in mES cells using a lower induction of Hoxb1. (A) Scheme of the experiment. Using the tet-ON/Hoxb1 mouse embryonic stem cell line (Gouti and Gavalas, 2008), Hoxb1 expression was induced by addition of 0.2µg/ml of doxycycline (DOX) from day 4 (D4) during mES cell differentiation into cardiac cells. (B) Quantification of beating embryoid bodies (EBs) relative to the total number of embryoid bodies. (C) Kinetics of expression for Hoxb1 and Myh6, Myh7, Mlc2v (or Myl2), Mlc2a during mES cell differentiation after induction of Hoxb1 expression (+ dox) or in control condition (no dox). Results are normalized for gene expression in undifferentiated mES cells (D0). Error bars indicate mean +/-SEM; n=2 experiments. Paired, one-sided t-test was performed based on relative transcript expression between control (no dox) and doxycycline treatment (+ dox). * indicates a significance level of p<0.05

**Figure 8 – Figure supplement 9:**
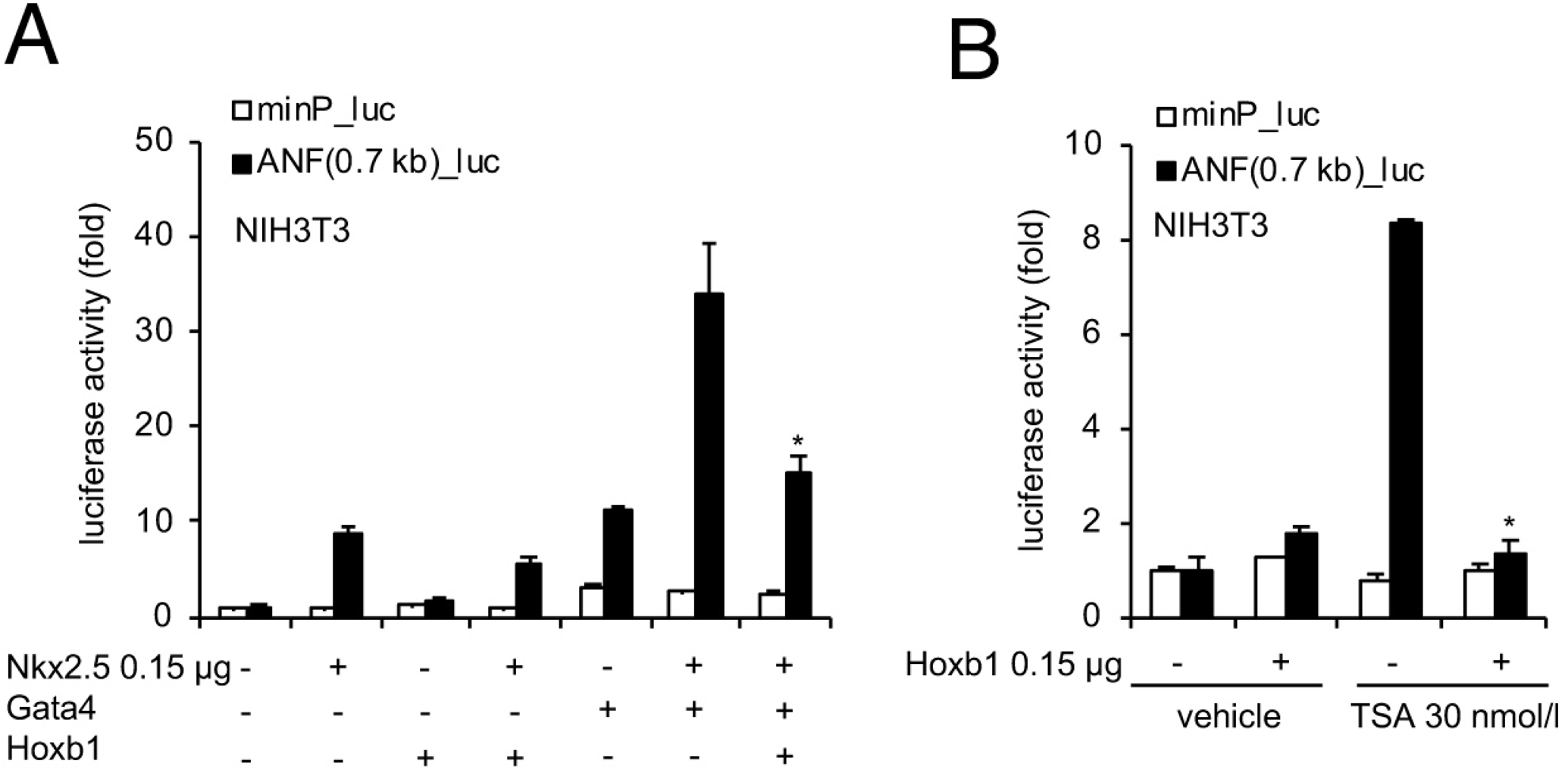
The 0.7-kb *Nppa* promoter is a functional target for Hoxb1. (A) Transient transfections were carried out with the 0.7-kb *Nppa* (ANF) promoter in NIH3T3 cells. Constructs were co-transfected with Nkx2-5, Gata4 and Hoxb1 expression vectors. Luciferase activity was determined and normalized as fold over the reporter alone (mean ± SEM, n=3, *p<0.05 for Nkx2-5 and Hoxb1 *versus* Nkx2-5, using ANOVA). (B) Luciferase reporter assays on the 0.7-kb *ANF* promoter in presence of TSA. NIH3T3 cells co-transfected with and without a Hoxb1 expression vector were treated in the absence or presence of 30 nmol/l TSA. Bars represent mean ± SEM (n=3). Statistical test was conducted using ANOVA (*p<0.05 for Hoxb1 and TSA treatment versus Hoxb1 or TSA treatment).

**Supplementary File 1.** Excel file containing the list of deregulated genes identified by RNA-seq analysis (sheet one, upregulated genes; sheet two, downregulated genes).

**Supplementary File 2.** Excel file containing the list of regions of open chromatin identified by ATAC-seq analysis (sheet one, upregulated genes; sheet two, downregulated genes).

